# Constrained proteome allocation affects coexistence in models of competitive microbial communities

**DOI:** 10.1101/2020.01.27.921478

**Authors:** Leonardo Pacciani-Mori, Samir Suweis, Amos Maritan, Andrea Giometto

**Author notes:** These authors contributed equally (Amos Maritan, Andrea Giometto). **Correspondence** Correspondence and requests for materials should be addressed to L. P.-M.

## Abstract

Microbial communities are ubiquitous and play crucial roles in many natural processes. Despite their importance for the environment, industry and human health, there are still many aspects of microbial community dynamics that we do not understand quantitatively. Recent experiments have shown that the metabolism of species in a community is intertwined with its composition, suggesting that properties at the intracellular level such as the allocation of cellular proteomic resources must be taken into account when describing microbial communities with a population dynamics approach. In this work we reconsider one of the theoretical frameworks most commonly used to model population dynamics in competitive ecosystems, MacArthur’s consumer-resource model, in light of experimental evidence showing how pro-teome allocation affects microbial growth. This new framework allows us to describe community dynamics at an intermediate level of complexity between classical consumer-resource models and biochemical models of microbial metabolism, accounting for temporally-varying proteome allocation subject to constraints on growth and protein synthesis in the presence of multiple resources, while preserving analytical insight into the dynamics of the system. We first show experimentally that proteome allocation needs to be accounted for to properly understand the dynamics of even the simplest microbial community, i.e. two bacterial strains competing for one common resource. We then study the model analytically and numerically to determine the conditions that allow multiple species to coexist in systems with arbitrary numbers of species and resources.

## Introduction

Microbes are among the most abundant life forms on Earth, in terms of biomass^1^. They are found in almost every habitat of our planet, and continue to surprise us with their ability to survive in places that were thought to be inhospitable and barren. For example, microbial communities have been found in the deep terrestrial subsurface^2,3^, and it has been estimated that the first five kilometers beneath the Earth’s surface could be habitable for them^4^. Because of their ubiquity, microbial communities play fundamental roles in countless natural processes of vital importance, from the digestion and overall health of their host organism^5^ to the regulation of bio-geochemical cycles^6,7^. Despite their importance, however, we still know very little about the fundamental mechanisms that regulate microbial communities, partly because we are only able to grow in the lab a very small fraction of all the microbes found in nature^8^, and partly because microbial communities are complex, non-linear systems^9^ whose dynamics is difficult to predict. For these reasons, scientists from many disciplines have long been fascinated by the challenging theoretical questions posed by the study of microbial communities’ structure and dynamics, and serious efforts are being made to understand how competition^10–12^ and metabolic interactions^13,14^ allow such systems to maintain the very high levels of biodiversity found in nature.

Recent experimental studies have shown that the structure and composition of microbial communities are tightly linked to the metabolism of the species that comprise them^15,16^ (e.g., communities with different taxonomic compositions can nevertheless exhibit the same metabolic functional structure^17,18^). We can therefore speculate that the ways with which microbes uptake and use different resources for growth and proliferation can affect the dynamics of an entire community. Resource uptake is constrained by the other functions that cells must perform to grow and proliferate, and the balance between such functions is governed by the allocation of the internal resources of the cell (e.g., the proteome, the set of proteins expressed by a cell) to different tasks. It is therefore important to understand how microbial community dynamics is influenced by the proteome allocation of its members, and new insights in this direction might help us make more powerful predictions of how microbial communities assemble and evolve^19,20^. However, accounting for the dynamics of metabolism and gene expression of each species in a microbial community explicitly (e.g., via community flux balance analysis^21^) can be very challenging, and the large dimensionality of the mathematical models that attempt to do so poses limits to our understanding of the dynamics of microbial communities and of the fundamental properties that affect species coexistence.

Scott *et al.*^22^ showed that, despite the complexity of bacterial metabolism, there are simple relationships that link the fraction of the proteome allocated for nutrient uptake and protein synthesis to the growth rate of bacteria grown in isolation, and that reducing these fractions by forcing cells to express a useless protein reduces their growth rate. Such relationships are very powerful because they describe how bacterial growth is influenced by proteome allocation and gene expression without requiring an explicit representation of the underlying molecular mechanisms. These relationships, which were also based on earlier observations by Schaechter *et al.*^23^ on how the ribosomal component of the proteome of a microbial species scales with the growth rate, have recently been applied in many different contexts^24^ and were instrumental in improving our knowledge of microbial metabolism, both experimentally^25^ and computationally^26^. However, as the experiments by Scott *et al.*^22^ were performed with single-species populations in exponential phase, it is still an open question if their approach can also be used to describe the population dynamics of different interacting microbial species competing for multiple resources.

In this work we fill this gap by linking the results by Scott *et al.*^22^ to one of the most widely adopted theoretical frameworks for modeling competitive ecosystems, MacArthur’s consumer-resource model^27–29^, and tailor it to describe the dynamics of microbial species (or strains) competing for one or more resources. MacArthur’s model describes how the population abundances of *N*_*S*_ species competing for a common pool of *N*_*R*_ resources change over time, and has been used in several recent studies^10–12,30–32^ to understand under which conditions multiple species can coexist while competing for few resources. We show that generalizing Scott *et al.*’s proteome-growth relationships and including them into a consumer-resource framework allows us to build a community dynamics model where all parameters can in principle be measured experimentally and have a precise biological interpretation. This “consumer-proteome-resource” model describes community dynamics at an intermediate level of complexity between classical consumer-resource models and biochemical models of microbial metabolism^21^. By adopting such an intermediate level of complexity and realism, we can take into account the dynamics of gene expression and microbial metabolism, while preserving analytical insights on the microbial community dynamics and identifying the key intracellular properties affecting species coexistence.

In the next section we describe our “consumer-proteome-resource” model for a general number of species/strains and resources. We then apply this model to the simplest implementation of an experimental microbial community, i.e. two *Escherichia coli* strains competing for glucose as the only carbon source. The experiment’s results highlight how the proposed theoretical framework accounting for proteome allocation allows us to understand species population dynamics in competitive communities. We finally study (both analytically and numerically) the “consumer-proteome-resource” model for communities composed of arbitrary numbers of species and resources to identify the conditions allowing the coexistence of multiple species in the community. A discussion section and some future perspectives conclude this work.

## Results

### The consumer-proteome-resource model

The phenomenological framework proposed by Scott *et al.*^22^ prescribes that the proteome of a single microbial species growing on a single resource can be minimally divided into three sectors: one dedicated to nutrient uptake and metabolism (the “P-sector”), one dedicated to ribosomal proteins responsible for biomass production and growth (the “R-sector”), and a third one dedicated to housekeeping functions (the “Q-sector”), which was shown to be incompressible^22^. Naming *φ*^*P*^, *φ*^*R*^ and *φ*^*Q*^ the proteome fractions corresponding to these sectors, we must have *φ*^*P*^ + *φ*^*R*^ + *φ*^*Q*^ = 1 (since all proteome fractions must sum to one), and Scott *et al.* have shown that *φ*^*P*^ and *φ*^*R*^ are linear functions of the species’ growth rate *g*, i.e:

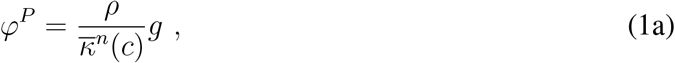

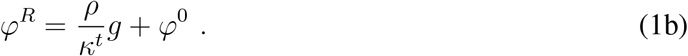

Here *ρ* is a conversion factor (equal to the ratio between the total mass of the ribosomal proteins and the total RNA mass of the cells) and 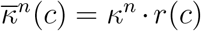, where *r*(*c*) = *c/*(*K* + *c*) is a Michaelis-Menten (or Monod) function, which encapsulates the dependence on the resource concentration *c*. *K* is the half-saturation constant of the resource and *κ*^*n*^ is the “nutritional capacity” of the (only) limiting resource. This parameter measures how much protein biomass is produced per unit ribosomal mass per unit time, and therefore depends on how much energy the resource contains and how efficiently the microbial species can metabolize it (see Supporting Online Material in Scott *et al.*^22^ for a molecular interpretation of *κ*^*n*^). The parameter *κ*^*t*^ is the “translational capacity” of the microbial species, measuring how much protein biomass is produced per unit ribosomal mass per unit time; it is therefore a measure of how fast the microbial species expresses its genome to synthesize proteins. Finally, *φ*^0^ is the incompressible core of *φ*^*R*^, representing the fact that ribosomal proteins are present in the cells also when microbes are not growing. All these parameters involve the ribosomal mass of the microbial species because the measurements by Scott *et al.*^22^ were done by assaying the RNA/protein ratio in exponentially growing *Escherichia coli*.

The way we generalize Scott *et al.*’s^22^ framework to a system with multiple species and resources is shown in Figure 1a: we assume that the proteome fraction 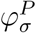 allocated by species *σ* to nutrient uptake and metabolism can be further sub-divided into smaller fractions 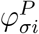, each one dedicated to one of the available resources. In other words, we call 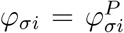 the proteome fraction allocated by species *σ* to the uptake and metabolization of resource *i*. With this choice, in order to ensure that the sum of all the proteome fractions is equal to one we must have:

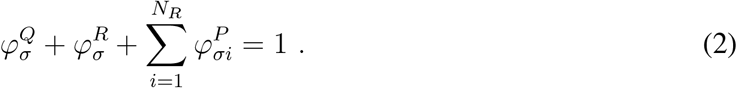

**Figure 1:**
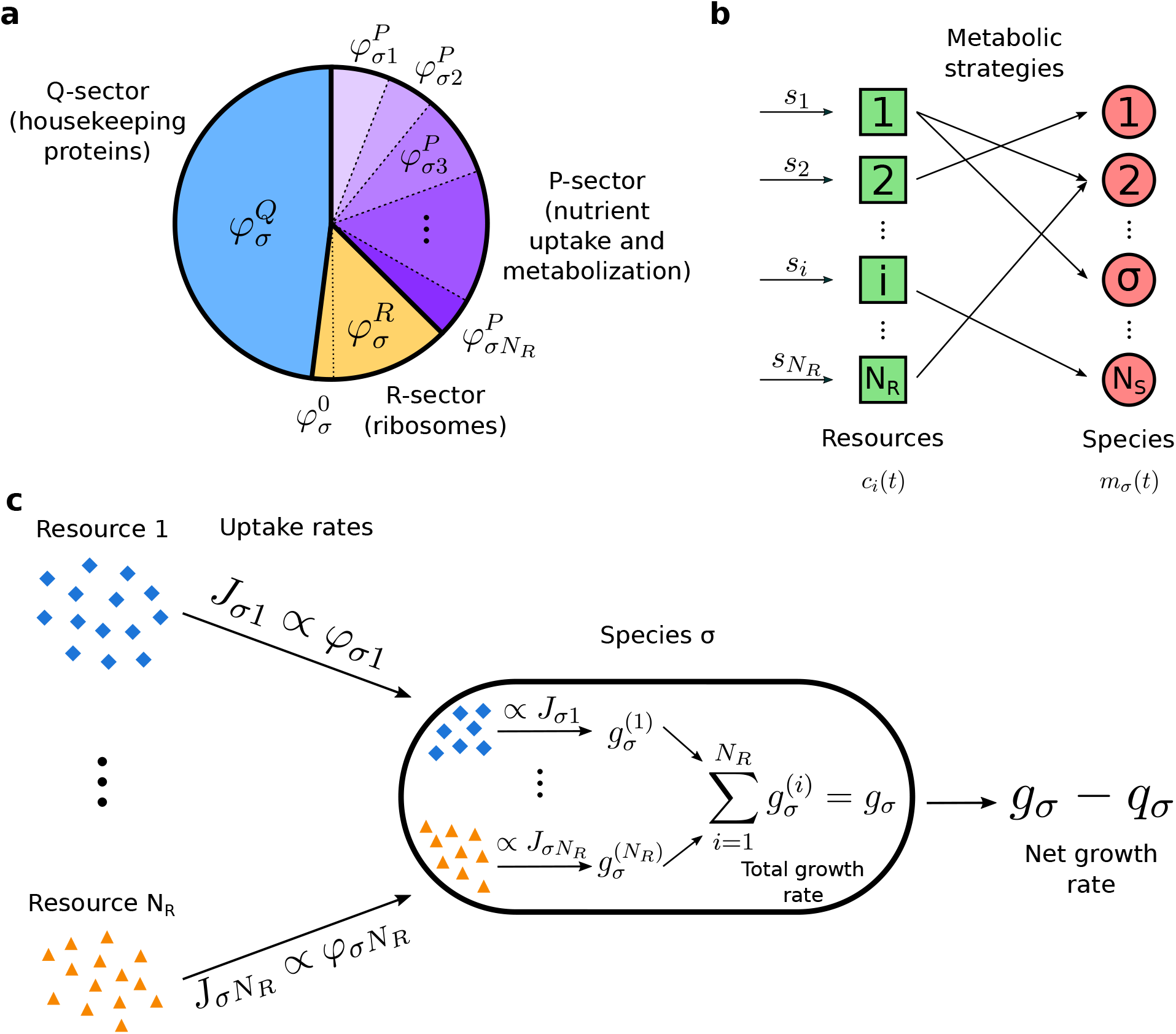
Assumptions made in building our model. **(a):** Generalization of the proteome subdivision introduced by Scott et al.^22^ to the case of *N*_*R*_ resources: we assume that the sector allocated by species *σ* for nutrient uptake and metabolization is subdivided into smaller fractions 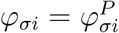, each one dedicated to a specific resource. **(b):** Schematic representation of a consumer-resource model with *N*_*R*_ resources and *N*_*S*_ species. In this framework, the concentrations *c*_*i*_ of the resources and the biomass densities *m*_*σ*_ of the species are described by systems of coupled differential equations. Resources are supplied with (constant) rates *s*_*i*_, and are uptaken by the species (arrows represent resource flows). The ways in which each species uptakes resources are encoded in the “metabolic strategies”. **(c):** Assumptions used to write the equations of our consumer-resource model with proteome allocation. Each species *σ* uptakes resource *i* with a rate *J*_*σi*_ proportional to the proteome fraction *φ*_*σi*_. Then, each resource contributes a growth term 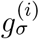 (proportional to the resource uptake rate) to the total growth rate of the cells. The net growth rate of species *σ* is the difference between this total growth rate and a maintenance cost *q*_*σ*_.

This constraint represents the finiteness of a microbial species’ proteome, i.e. the fact that each species in a community has a limited “proteomic budget” that can be spent for all the necessary biological functions: for example, if more proteins need to be produced for metabolizing complex substrates (i.e., if the nutrient fraction 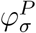 increases), then a smaller part of the proteome will be available for biomass production (i.e., the ribosomal fraction 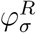 decreases). In order to achieve optimal growth, microbial species must balance this trade-off^22^.

In Figure 1b we show a schematic representation of the “classic” consumer-resource model. Within this framework, a community is a set of *N*_*S*_ species that can only uptake some (or all) of the *N*_*R*_ available resources. The species’ growth is determined by the types and the amout of resources they uptake, and is also regulated by a “maintenance cost” (representing the fact that species need to uptake a minimum amount of resources in order to survive). The resources, on the other hand, can be thought of as organic substrates that are supplied to the system with given (constant) rates *s*_*i*_. The model describes explicitly the dynamics of both species and resources, and the ways in which the species uptake the available substrates are encoded in parameters that in the literature are called “metabolic strategies” or “resource preferences”. Therefore, in the consumer-resource framework the interactions between species are indirect and mediated by the abundance of resources, and other types of direct inter-specific interactions (like the exchange of useful metabolic byproducts) are not included.

Figure 1c depicts schematically the assumptions underlying our consumer-proteome-resource model. Each species *σ* uptakes resource *i* with a rate *J*_*σi*_ that is proportional to the proteome fraction *φ*_*σi*_. Then, resource *i* is converted into a growth term 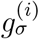 proportionally to the uptake rate *J*_*σi*_. For our purposes, we assume that all resources in the system are substitutable, so that they can be used interchangeably and we can write the total growth rate *g*_*σ*_ of a given species as the sum of all the terms 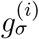. Eventually (see Materials and Methods for all the detailed computations) we obtain a mathematical model that has the traditional structure of a consumer-resource model, but with the added merit of describing population dynamics using parameters and variables that have a precise biological meaning at the intracellular scale of the system and that can in principle be measured experimentally^22^. For example, “metabolic strategies” in our framework correspond to the proteome fractions *φ*_*σi*_. Additionally, our model also includes the proteome finiteness constraint given by Eq (2). The final expression of this constraint in our equations is significantly different from similar constraints that have been studied in the consumer-resource framework^31,33^, and all the results that we will show are ultimately a repercussion of this finiteness of the microbial proteome, as encoded in our model.

The most important consequence of the proteome finiteness constraint is that it implies that the proteome fractions *φ*_*σi*_ *cannot* be fixed parameters, but must change over time and therefore must be dynamical variables (see Materials and Methods). It is thus necessary to introduce some form of dynamics on the proteome fractions that each species allocates for nutrient uptake and metabolization. Our approach is to require that all *φ*_*σi*_ evolve dynamically with a characteristic timescale in order to maximize the instantaneous growth rate^33^ of species *σ* in what we call an “adaptive process”, while we ensure that the proteome finiteness constraint is satisfied at all times. The mathematical details are discussed in the Materials and Methods section.

### Application of the model to an experimental competitive microbial community

We have applied our consumer-proteome-resource model to the simplest case of a competitive community, i.e. two species competing for one common resource. We competed experimentally two strains of *E. coli* grown in a liquid minimal medium with glucose as the sole carbon source, transferring a fraction of the community to fresh medium daily and measuring the relative abundance of the two strains at each transfer (see Materials and Methods). The two strains had the same genetic background and expressed constitutively from their genome two different fluorescent proteins, which allowed us to measure their relative abundance via flow-cytometry. Additionally, we introduced in strain *σ* = 1 a plasmid containing a red fluorescent protein (RFP) whose expression could be controlled by adding Isopropyl *β*-D-1-thiogalactopyranoside (IPTG, a molecular mimic of allolactose that cannot be metabolized by *E. coli*) to the medium. Thus, by varying the concentration of IPTG in the medium we could vary the proteome allocation of strain 1 by forcing it to produce a useless protein. The experiment is sketched in Figure 2.

**Figure 2:**
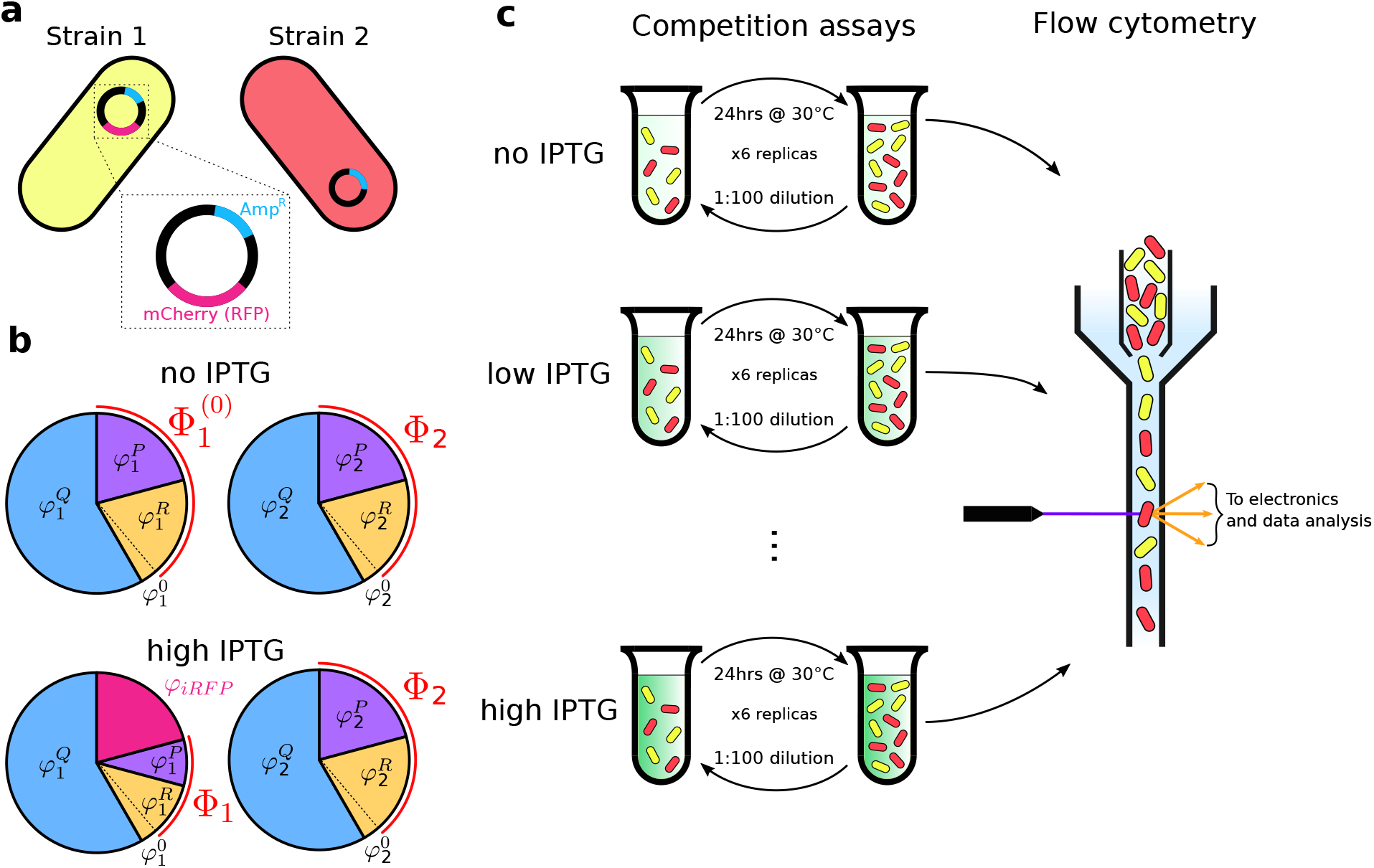
Schematic representation of the experiment. **(a):** Two *E. coli* strains were used: strain 1 constitutively expresses a yellow fluorescent protein (mVenus) and carries a plasmid with the ampicillin resistance cassette (cyan Amp^*R*^ in the plasmid magnification) and a red fluorescent protein (RFP), mCherry (magenta), under the control of the *trc* promoter, an hybrid of the *trp* and *lac* promoters. Strain 2 constitutively expresses a red fluorescent protein (mKate2Hyb) and carries a plasmid with the ampicillin resistance cassette. **(b):** Proteome allocation of the two strains at different concentration of IPTG in the medium. When strain 1 grows in the presence of IPTG, a fraction *φ*_*iRFP*_ of the strain’s proteome is allocated for the expression of the RFP mCherry, thus reducing the fraction Φ_1_ allocated for metabolism and growth. The proteome allocation of strain 2, instead, is not affected by the presence of IPTG. **(c):** The two strains were co-cultured in minimal medium at different IPTG concentrations, they were diluted daily into fresh medium and their relative abundance was measured at every transfer via flow-cytometry.

Applying our model to such a simple community, and introducing assumptions consistent with our experimental conditions (e.g., the fact that the strains are grown in medium-rich conditions, and that they share the same genetic background), our model predicts that the selective advantage 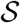 of strain 1 over strain 2 (a measure for the difference in fitness between the two strains) is:

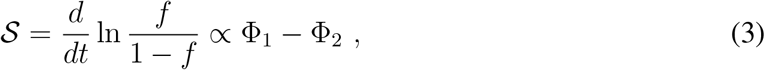

where *f* is the relative abundance (or frequency) of strain 1, and Φ_*σ*_ is the total proteome fraction allocated by species *σ* for metabolism and growth (see the Materials and Methods section for all the details). In other words, the selective advantage of strain 1 over strain 2 is proportional to the difference between these two proteome fractions.

According to Eq (3), the ratio between the relative abundances of the two strains decreases or grows exponentially with time, depending on the sign of Φ_1_ − Φ_2_, which then sets the outcome of competition: for example, if Φ_2_ *>* Φ_1_ (i.e., strain 2 allocates a larger fraction of its proteome to metabolism and biomass production than strain 1) then 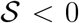 and strain 2 outcompetes strain Coexistence between the two strains is possible uniquely when Φ_1_ = Φ_2_ and thus 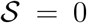. The system therefore exhibits two regimes where only one of the two strains survives (competitive exclusion), separated by the coexistence point Φ_1_ = Φ_2_. Eq (3) thus connects a well known concept of population genetics, the selective advantage in exponentially growing populations, with the differential proteome allocation Φ_1_ − Φ_2_ between microbial strains. According to this model, a 1% difference in proteome allocation for growth between two strains (i.e., Φ_1_ − Φ_2_ = 1%) corresponds to a selection coefficient of 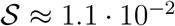 1/h (see Materials and Methods for details on this estimation). Considering now that we can force strain 1 to produce a useless RFP, we can investigate how changing its proteome allocation affects the outcome of competition. Indicating with *φ*_*iRFP*_ the fraction of proteome allocated by strain 1 to the synthesis of the IPTG-inducible RFP, the proteome fraction allocated for nutrient uptake and growth is 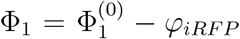 (with 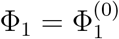 in the absence of IPTG), and we find that the selective advantage 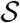 is predicted to decay linearly with *φ*_*iRFP*_ as 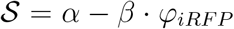 with *α* and *β* positive constants (see the Materials and Methods section for all details and the explicit expression of 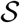 in this case).

As shown in Figure 3a (magenta data points), the experimental selective advantage 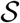 decreases linearly with the production rate of the IPTG-inducible RFP of strain 1 over a broad range or RFP production rates (the mean cell’s fluorescence measured after 8 h at 105 *μ*M IPTG is 22 times higher than at 0 *μ*M IPTG, Figure 3d), which are proportional to *φ*_*iRFP*_. In the absence of IPTG and at low concentrations of it, strain 1 outcompetes strain 2 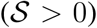. At an IPTG concentration of approximately 30 *μ*M, the two strains coexist by maintaining a stable relative fraction. At IPTG concentrations larger than 30 *μ*M, strain 1 is outcompeted by strain 2 (i.e., 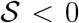). By manipulating the proteome allocation of strain 1 we are thus able to explore the expected regimes of the system. Figure 3a also shows that strain 1 has a fitness advantage over strain 2 in the absence of IPTG, since 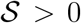 at low protein production rates. In our theoretical framework, such an advantage implies that 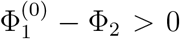. It is also possible to use our data to estimate the ratio Φ_1_/Φ_2_ at different protein production rates, shown in Figure 3c (see Materials and Methods for details). This ratio is approximately 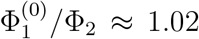 for low protein production rates and then decays linearly up to Φ_1_/Φ_2_ ≈ 0.98.

**Figure 3:**
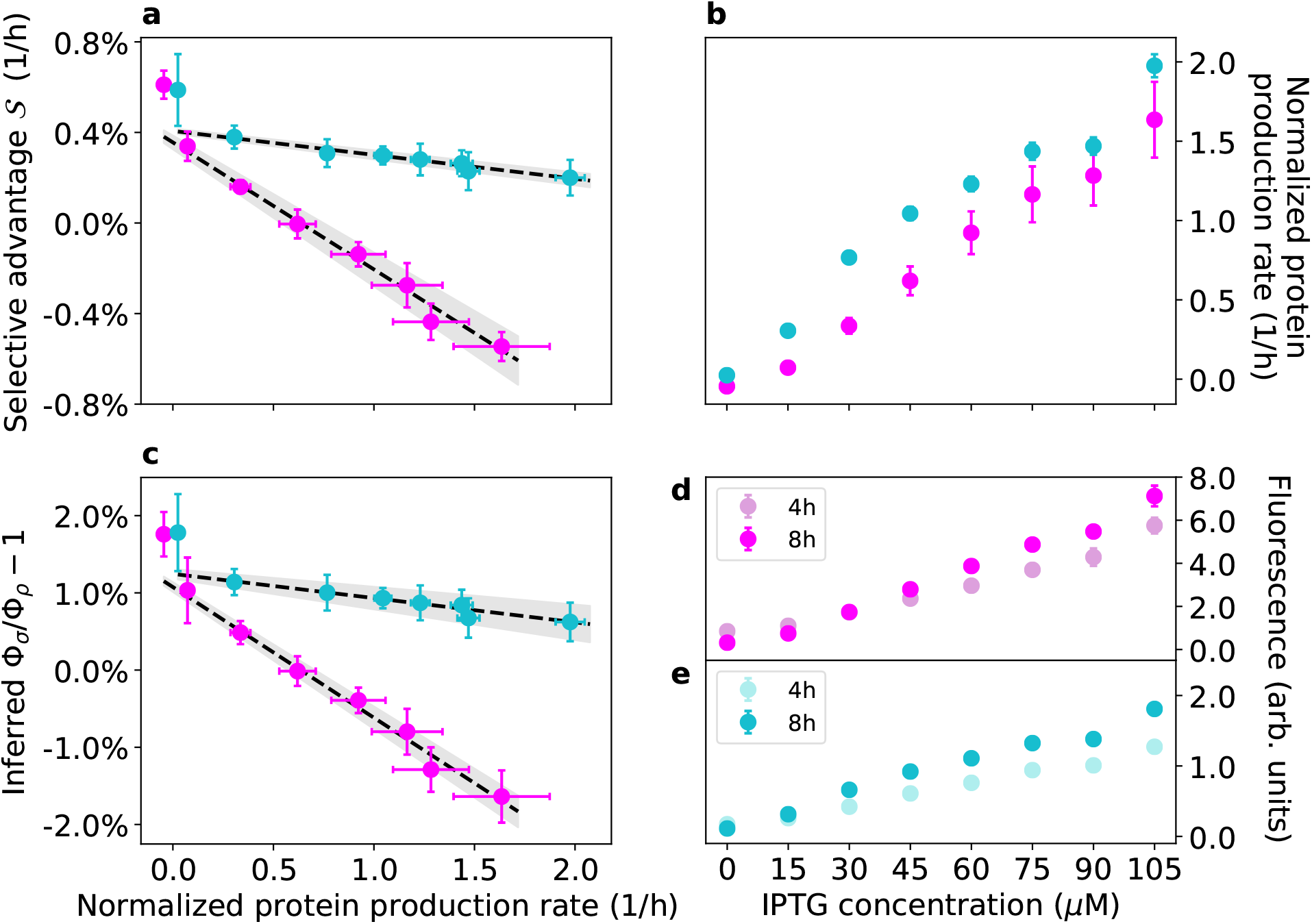
Experimental results. Magenta points represent data from experiments with strains 1 and Cyan points represent data from experiments with strains 3 and 4, where strain 3 expresses constitutively mKate2Hyb and the IPTG-inducible Venus yellow fluorescent protein (YFP) and strain 4 expresses mVenus constitutively (see Materials and Methods and Figure S.1). Error bars represent two standard deviations. Note that the normalized protein production rates are not directly comparable across magenta and cyan points (see Materials and Methods). **a:** The experimental selection coefficient 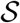 (y axis) decreases linearly with the normalized production rate of the inducible protein, measured as the temporal variation of the mean cell fluorescence signal at different concentrations of the inducer IPTG, accounting for dilution of such protein via cell division (see Materials and Methods). The grey band represent the 68% confidence interval of the linear fit. The time series of ln[*f*/(1 − *f*)] for the two experiments are reported in Figures S.2 and S.3. **b:** Induced protein production rates as functions of IPTG concentration (see Materials and Methods). **c:** Inferred values of the ratios Φ_1_/Φ_2_ and Φ_3_/Φ_4_ (minus one) as functions of the induced protein production rate (normalized). Also shown are the linear fits of the data with their 68% confidence interval. **d:** Mean (induced) red fluorescence of strain 1 at 4h and 8h after inoculation in the conditions used for our experiment (see Materials and Methods). **e:** Mean (induced) yellow fluorescence of strain 3 at 4h and 8h after inoculation in the conditions used for our experiment (see Materials and Methods).

Figure 3 also shows the results of a second experiment performed with two different strains (cyan data points). These strains have different fluorescent protein combinations with respect to strains 1 and 2 (see Materials and Methods and Figure S.1): strain 3 expresses constitutively a red fluorescent protein (mKate2Hyb) and carries a plasmid with an IPTG-inducible yellow fluorescent protein (Venus YFP), while strain 4 expresses constitutively the yellow fluorescent protein mVenus (see Materials and Methods). Also in these independent sets of experiments, the selective advantage decreases linearly as the protein production rate is increased over a broad range (the mean cell’s fluorescence measured after 8 h at 105 *μ*M IPTG is 16 times higher than at 0 *μ*M IPTG, Figure 3e). In the first set of experiments (magenta points in Figure 3), and to a lesser degree in the second set of experiments (cyan points), the data points at the lowest production rate (i.e., at 0 *μ*M IPTG) appear to deviate from the linear trend, and so the fits in Figure 3a-c were calculated by excluding those data points (including those points in the fit doesn’t affect the results, see Figure S.20). The flow-cytometry data suggest that the average fluorescent intensity of strain 1 from the induced RFP decreased over the course of the experiment at 0 *μ*M IPTG, which may partly explain the deviation of the first magenta point in Figure 3a from the linear trend via a reduction in protein production rate throughout the experiment at 0 *μ*M IPTG. Another factor that may cause deviations from a linear trend is an increased gene-expression heterogeneity between cells in the absence of IPTG, a well-known property of the lac operon whose constituent parts we have used in our genetic constructs^34^, which might confer heterogeneous growth rates to different cells in the population. Please note that the normalized protein production rates of the two sets of experiments (magenta and cyan data points) are not directly comparable (see Materials and Methods).

### Coexistence of multiple species

An analytical and numerical analysis of our consumer-proteome-resource model in the general case of multiple species and multiple resources can provide some insights into the conditions needed for species coexistence. In particular, looking for stationary solutions of our model where all species have non-null biomass densities yields two necessary conditions (see the Materials and Methods section for all detailed expressions and computations). The first one is that the maintenance cost *q*_*σ*_ of species *σ* must be proportional to the total proteome fraction allocated for metabolism and growth, i.e. *q*_*σ*_ ∝ Φ_*σ*_, with a species-dependent proportionality constant. This requirement is biologically reasonable, since allocating a larger fraction of the proteome to such functions requires additional energy to synthesize the necessary proteins.

The second condition resembles similar ones that have been shown to hold for consumer-resource models with metabolic trade-offs^31^, and can be interpreted graphically as follows (see Materials and Methods for all the mathematical details). A system with *N*_*R*_ resources can be represented on an (*N*_*R*_ − 1)–dimensional simplex, where each vertex corresponds to one of the available resources; considering for example the case *N*_*R*_ = 3, the system can be represented on a triangle (i.e., a bi-dimensional simplex) as shown in Figure 4. On this simplex we can draw the vectors 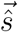 and 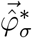, whose components are appropriately rescaled versions of (respectively) the resource supply rates *s*_*i*_ and the stationary proteome fractions 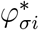 (i.e., the values of the proteome fractions once the species’ populations have reached a stationary state). The second condition for species coexistence is therefore that 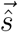 must belong to the *convex hull* of the vectors 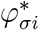, as shown in Figure 4. Notice that this condition involves the *stationary* proteome fractions 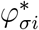. Since the model is highly non-linear, it is impossible to predict *a priori* these stationary values in general.

**Figure 4:**
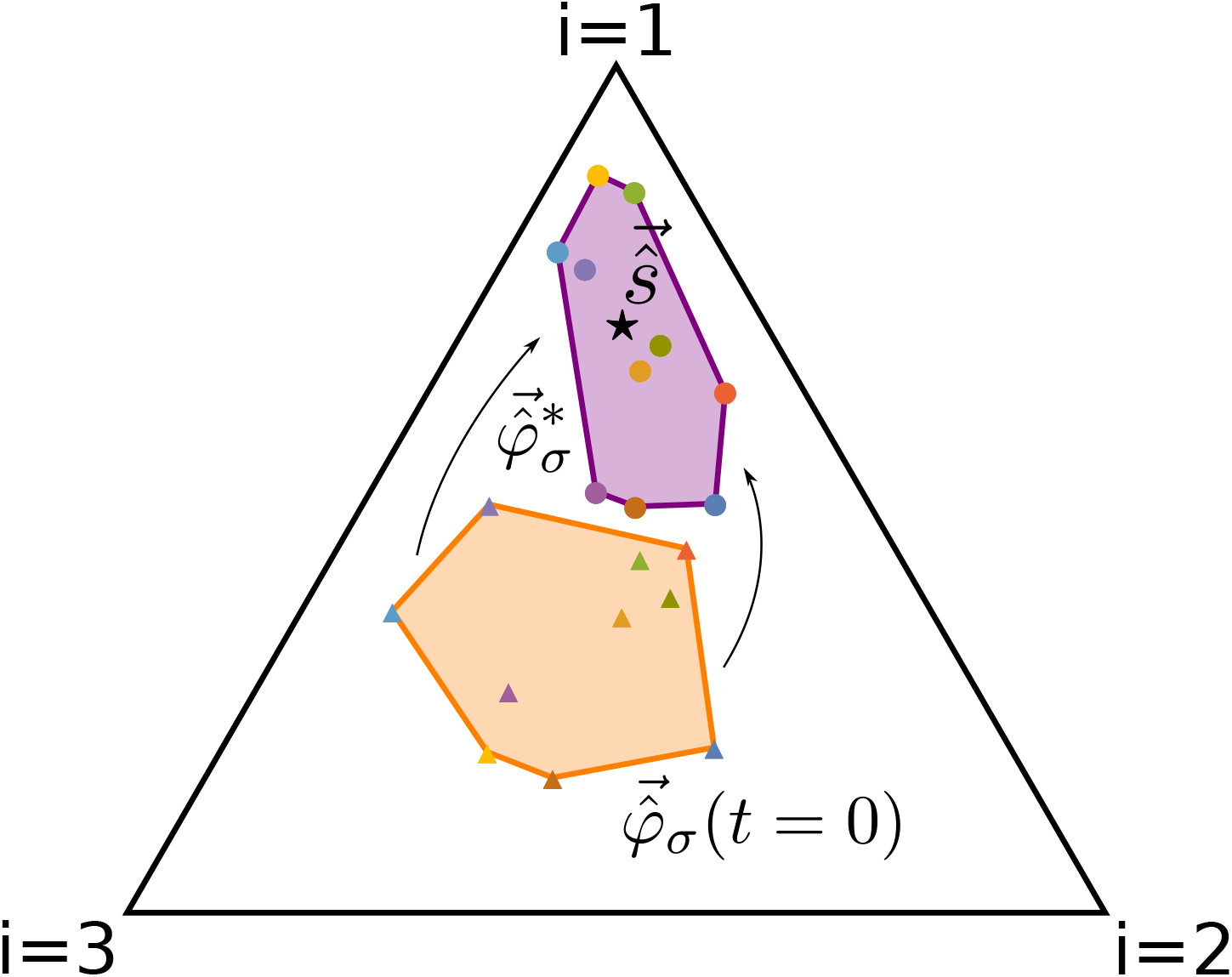
Graphical representation of the second condition necessary for coexistence. Here we consider a system with *N*_*S*_ = 10 species and *N*_*R*_ = 3 resources (for ease of representation). In this case, the system can be represented on a bi-dimensional simplex (i.e., a triangle) where each vertex corresponds to one of the available resources. On this simplex, we can draw the rescaled nutrient supply rate vector 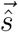 (black star) and the rescaled initial proteome fractions 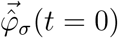 allocated by the species to the uptake and metabolism of the resources (colored triangles); their convex hull is drawn in orange. We have also drawn the stationary values 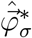 of the proteome fractions (colored circles), and their convex hull is drawn in purple. In this representation, if 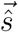 lies on one on the sides of the simplex, it means that only two of the available resources are being externally supplied to the system, and analogously if one of the 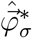 lies on one of the sides of the simplex, it means that its corresponding species is uptaking and metabolizing only two of the available resources. In general, the positions of 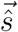 and 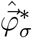 depend on the relative ratios with which the resources are supplied or uptaken by the species. Using this representation, our model prescribes that coexistence will be possible if 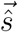 lies inside the convex hull of the stationary fractions 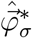 as shown in this case.

However, by exploring different regions of the parameter space it is possible to make predictions. In fact, the parameters that are relevant in this sense are the following: the ratios 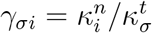 between the nutritional and translational capacities and the characteristic timescales *τ_σ_* of the adaptive process that maximizes the growth rate *g*_*σ*_ in the dynamics of *φ*_*σi*_ (see the Materials and Methods section for details). These timescales *τ*_*σ*_ measure how fast the dynamics of the proteome fractions *φ*_*σi*_ vary: the smaller *τ*_*σ*_, the faster species *σ* can switch between different resources. Biologically speaking, this parameter can be thought of as a measure of how “fast” the regulatory mechanisms of a microbial species can respond to changes in the availability of resources.

The first regime that we have explored is *τ*_*σ*_ ≫ 1 and *γ*_*σi*_ ~ 0, or in other words: the species’ adaptive process is very slow (i.e., they respond very slowly to changes in resource abundance) and the nutritional capacity is much smaller than the translational capacity (which happens, for example, when species are grown in very low quality nutrients). In this case the model predicts that the stationary values 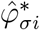 of the proteome fractions allocated by the species to nutrient uptake and metabolization change negligibly, and therefore the rescaled nutrient supply rate vector 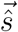 must lie in the convex hull of the rescaled *initial* proteome fractions 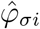, as shown in Figure 5.

**Figure 5:**
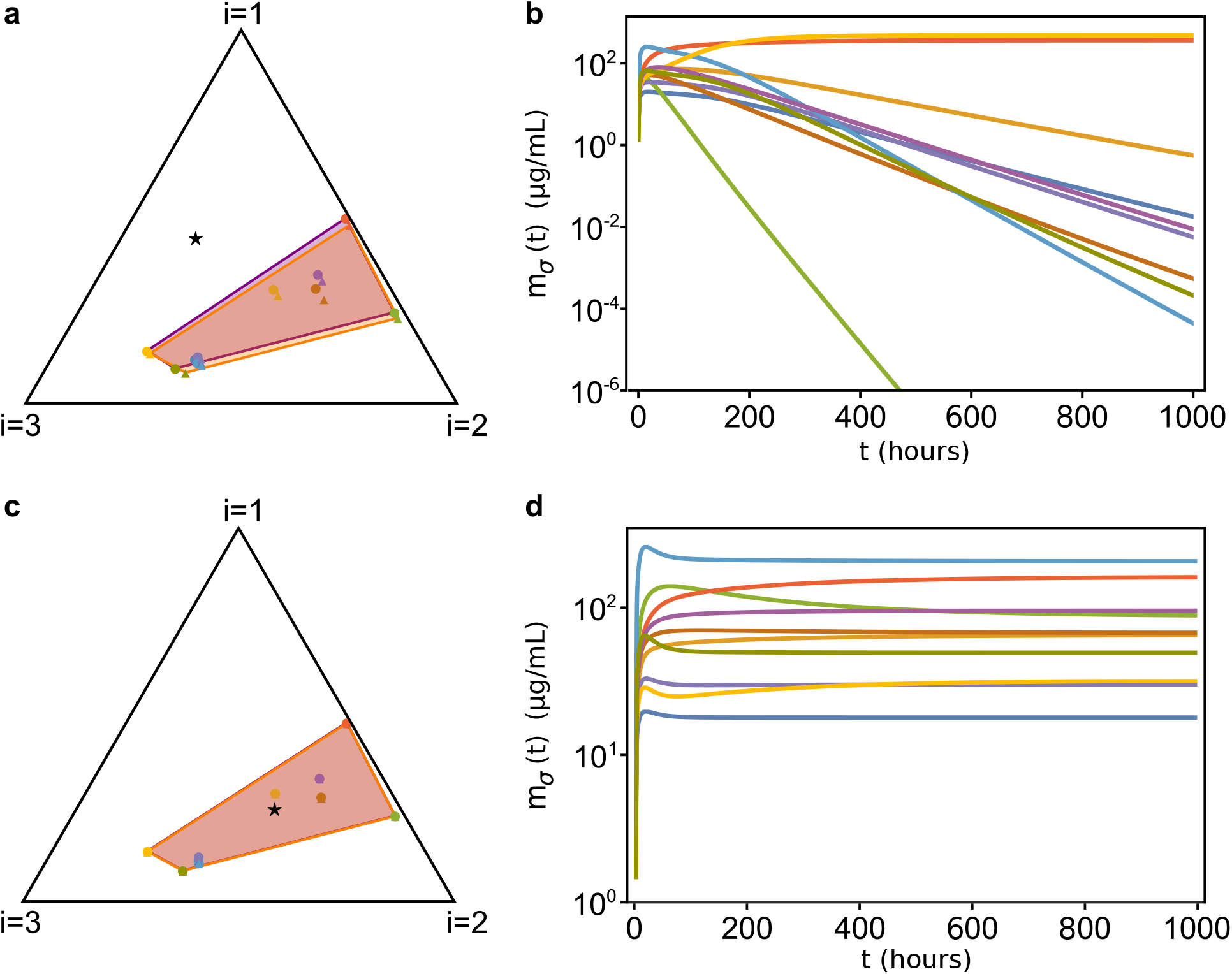
Temporal evolution of our consumer-proteome-resource model when *τ*_*σ*_ ≫ 1 and *γ*_*σi*_ ~ 0.1. **(a):** Initial conditions for the 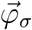 of a system with 10 species and 3 resources. We are using the graphical representation^31^ shown in Figure 4: the black triangle is the simplex to which the 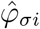 (colored dots) and the 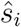 (black star) belong. The initial 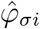 are represented as colored triangles, and their convex hull is colored in orange, while 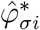 are represented as circles of the same colors, and their convex hull is in purple. With good approximation, 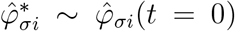. **(b):** Time evolution of the species’ biomasses *m*_*σ*_ relative to the case shown in (a). Since 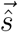 lies outside of the convex hull of the 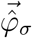, most species go extinct. **(c):** Same as in (a), but with 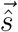 belonging to the convex hull of the 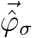. **(d):** Population dynamics of the system corresponding to the case shown in (c). In this case all species coexist. The parameters and the initial conditions were drawn from random distributions (see SI for more information). All parameters other than 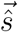 are identical in the four panels).

In the second regime we still have *τ*_*σ*_ ≫ 1, but now *γ*_*σi*_ ≳ 1 (i.e., the ratios between the nutritional and translational capacities have slightly larger values). In this case the dynamics of 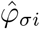 allows the proteome fractions to move inside the simplex. Therefore, the system can reach stationary states where all species coexist even when 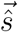 is not necessarily close to the convex hull of the initial 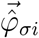. On the other hand, we observed that if 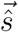 is too far away from the initial convex hull of 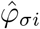 there might be extinctions. However, if 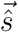 lies at an intermediate distance between these two cases, the system can reach diverse stationary states only if the resource supply rates *s*_*i*_ are sufficiently large. For example, multiplying each resource supply rate by a factor *x* > 1, i.e. *s*_*i*_ → *xs*_*i*_ (this rescaling leaves 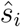 unchanged, see Materials and Methods), we observe a transition between two different states of the system for increasing values of *x*: when *x* ~ 1, only one or a few strains survive, whereas for larger *x* the stationary biomass densities 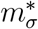 of the other species increase until all of them coexist. Figure 6 shows an example of such transition. This phenomenon occurs only when 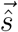 lies in specific areas of the simplex, whose shape and position can be determined numerically, but depend on the particular values of the model parameters used.

**Figure 6:**
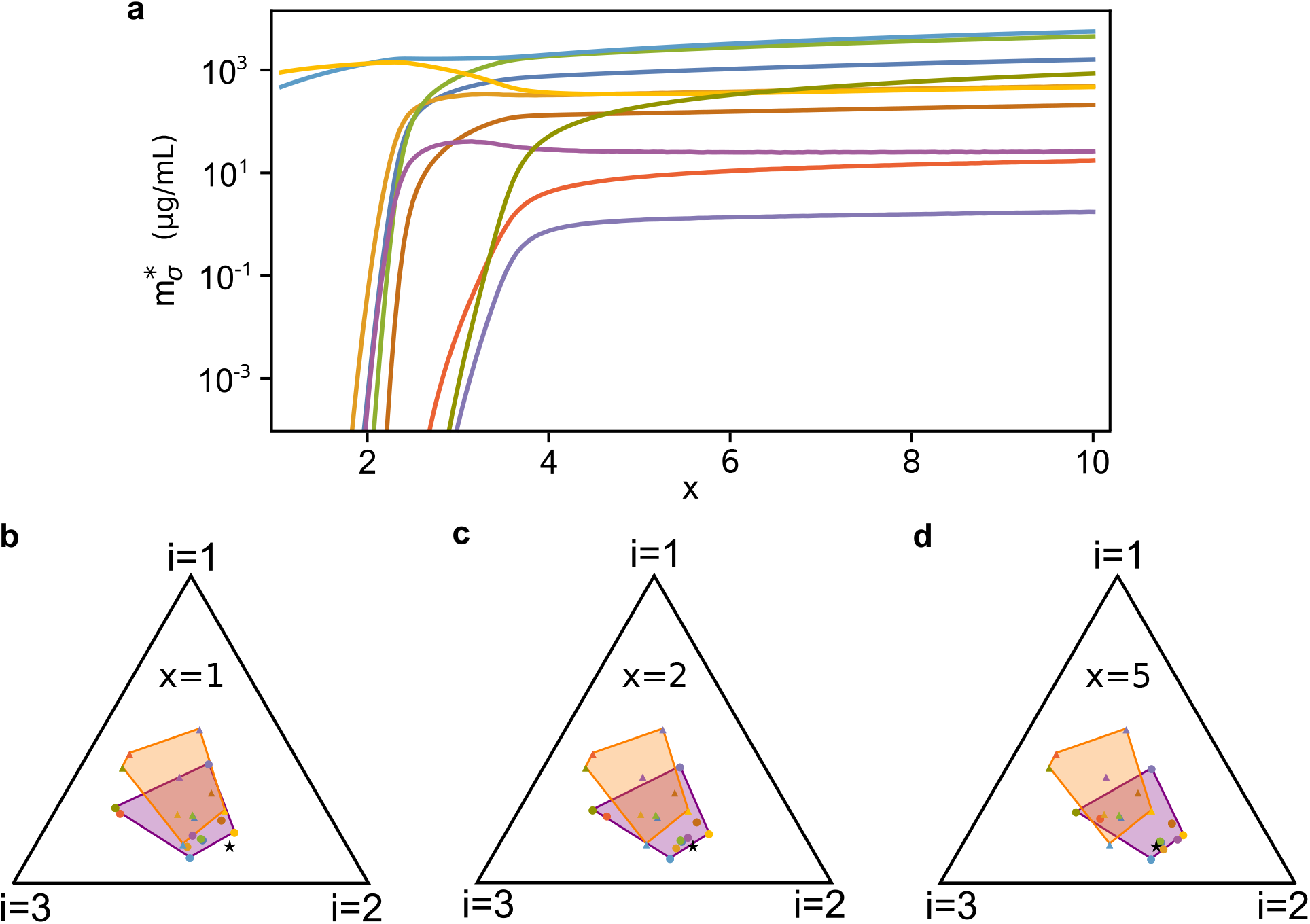
Species coexistence as a function of the rescaled resource supply rate 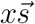 (with *x* > 1). As for Figure 5, the 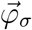 evolve according to our model with *τ*_*σ*_ ≫ 1, *γ*_*σi*_ ≳ 1, *N*_*S*_ = 10 and *N*_*R*_ = 3. Here, 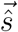 was drawn randomly outside the convex hull of the initial 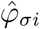 (same 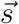 for all panels) and we varied *x* > 1. **(a):** Stationary values of the species’ biomasses for different values of *x*. When *x* ⋍ 1 the system is in an oligodominant phase where only one or a few species survive, but as *x* grows larger the system shifts to a diverse phase in which all species coexist. Notice that the relative ratios of the stationary abundances 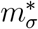 are not constant as *x* grows. **(b-d):** Initial (orange) and stationary (purple) convex hull of the rescaled proteome fractions 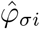 for different values of *x*. For small *x*, the resource supply (black star) is not large enough to allow the 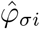 to move so that coexistence condition is satisfied. Increasing *x* (d), this becomes possible and thus all species coexist. The parameters and the initial conditions were drawn from random distributions (see SI for more information). All parameters other than 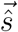 and the initial conditions *m*_*σ*_(0) and *c*_*i*_(0) are identical in the four panels).

In this same regime, if *γ*_*σi*_ assume increasingly large values (which happens for example, if the species are grown in nutrients with increasingly higher qualities) coexistence will be possible even if 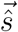 lies at increasingly large distances from the convex hull of the initial 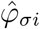.

Finally, the last regime that we explored is *τ* ≲ 1, i.e. the adaptive process that maximizes the species’ growth rates is fast. In this case, the smaller the timescales *τ*_*σ*_ are, the faster the proteome fractions *φ*_*σi*_ will reach their stationary values, and coexistence will always be possible. In other words, if the timescales *τ*_*σ*_ have sufficiently small values, all species will be able to coexist independently of the initial values of the proteome fractions *φ*_*σi*_ and of the resource supply rates *s*_*i*_. However, as the *τ*_*σ*_s grow, fewer and fewer species will be able to coexist. This can be shown by multiplying *τ*_*σ*_ by a factor *y* > 0: Figure 7 shows how the species’ stationary biomasses change as *y* increases, and we can see that as species adaptation becomes slower (i.e., for larger *y*), fewer and fewer species survive in the community.

**Figure 7:**
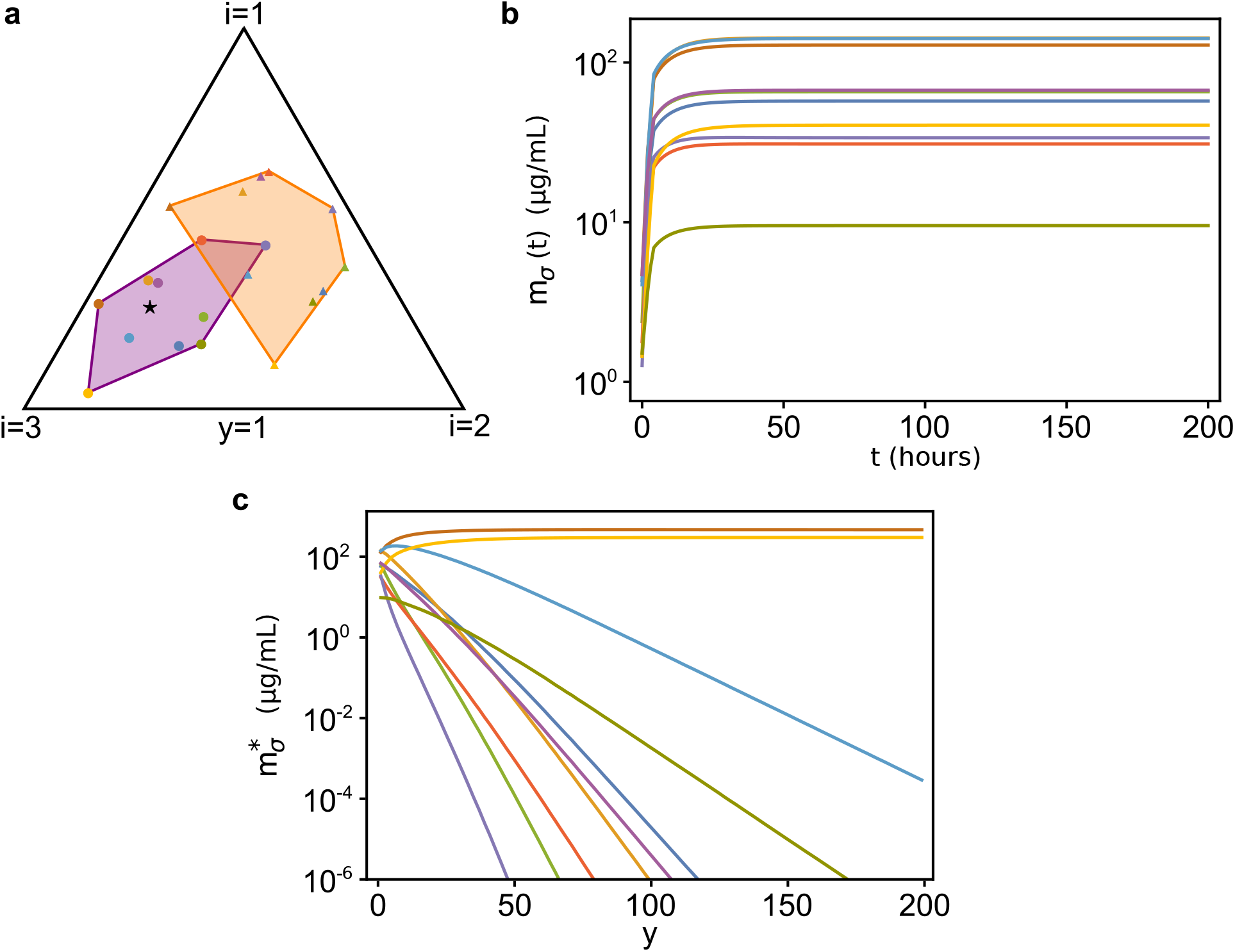
Temporal evolution of our consumer-proteome-resource model with *τ*_*σ*_ ≲ 1, *N*_*S*_ = 10 and *N*_*R*_ = 3. **(a):** Initial (orange) and stationary (purple) convex hull of the rescaled proteome fractions 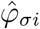. **(b):** Population dynamics of the system represented in (a). **(c):** Stationary values of the species’ biomasses as function of *y*, where in this case we have used *yτ*_*σ*_ with *y* > 1 as the timescales over which the 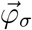 evolve (and all other parameters have been kept equal to the cases shown in (a) and (b)). As we can see, as *y* increases the system shifts from a diverse stationary state to one where only few species survive. The parameters and the initial conditions were drawn from random distributions (see SI for more information). All parameters other than *y* are identical in the three panels).

## Discussion

Motivated by our experiment that shows how varying proteome allocation can have strong effects on the dynamics of a very simple microbial community, we have formulated a consumer-resource model that generalizes and incorporates the phenomenological laws discovered by Scott *et al.*^22^. In this way we have related microbial growth to proteome allocation in competitive communities and have investigated the conditions that lead to species coexistence in the presence of multiple resources.

Our model describes the population dynamics of a purely competitive microbial community, i.e. an ensemble of species that compete directly for the same pool of resources. Our main contribution is introducing a physiological, experimentally-validated constraint on the amount of resources that cells can devote to growth and metabolism in consumer-resource models with temporally-varying nutrient uptake rates, differing from previous works^31,33^ that considered phenomenological constraints not based on direct experimental measurements. Introducing the right constraint in such models is particularly important, because the exact conditions that allow species coexistence depend on the specific form of the constraint (see Materials and Methods).

We have shown that the model can make meaningful predictions when applied to a system of two bacterial strains competing for one common resource. Furthermore, when larger communities with multiple species and resources are considered, the model predicts that high levels of biodiversity can be achieved only if certain conditions apply. In particular, we find that coexistence is possible when two conditions apply. The first one is that the maintenance cost must be proportional to the total proteome fraction allocated by the species to metabolism and growth, i.e. *q*_*σ*_ ∝ Φ_*σ*_. The second one can be interpreted graphically as described in the Results section, and summarized as follows: *i)* if the timescales *τ*_*σ*_ over which the species shift between different resources are large (i.e., *τ*_*σ*_ ≫ 1) and if the quality of the resources is low, coexistence will be possible only if the resource supply rates have particular values (i.e., the rescaled nutrient supply rate vector 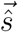 belongs to the convex hull of 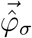); *ii)* if again *τ*_*σ*_ ≫ 1, but the resources are of higher quality, coexistence becomes possible (and in some cases the magnitude of the resource supply rates must be large enough); *iii)* if the resources’ quality is higher, coexistence is favored; and *iv)* coexistence is favored for smaller values of the timescales *τ*_*σ*_.

The dynamics of microbial communities has traditionally been studied at the ecological level by using models of population dynamics describing how the population abundances of different species in the community change over time as the result of competition for resources. While this approach is undoubtedly useful and effective, it often cannot describe the system at a level of detail necessary to make predictions from measurable quantities. In fact, it is becoming increasingly clear that the structure and dynamics of microbial communities are affected by the metabolic activity of the species that comprise them^15–18^. As shown here, mathematical models of community dynamics that take explicitly into account how different species allocate their proteome to regulate nutrient uptake can provide new insights into the link between the ecological properties of microbial communities, i.e. population dynamics and species coexistence, and their intracellular ones, i.e. metabolism and gene expression^20^.

Direct competition for resources is only one of the many known interactions that can take place between microbial species: exchange of metabolic byproducts^14^, production of toxins^13^ and environmental conditioning^35^ are only a few of the ways in which we know microbes interact within a community. Each of these processes provide both growth benefits and proteomic costs to microbial species, and can in principle be included in our framework by appropriately taking into account how they affect proteome allocation and species’ fitness. With our framework it would therefore be possible to make quantitative predictions involving such phenomena, and testing them against experimental data.

## Materials and Methods

### The consumer-proteome-resource equations

The general structure of a consumer-resource model is:

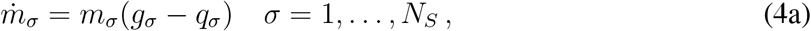

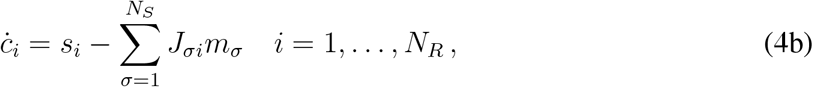

where *m*_*σ*_ is the biomass density of species *σ* and *g*_*σ*_ is its growth rate. The parameter *q*_*σ*_ is a maintenance cost, due to the fact that each species requires a minimum amount of energy per unit time to survive without growing. Finally, *c*_*i*_ is the density of resource *i*, *s*_*i*_ is the (constant) resource supply rate, and *J*_*σi*_ is the rate at which species *σ* uptakes resource *i* per unit biomass. To write these equations explicitly, we introduce the following assumptions: *i)* the uptake rate *J*_*σi*_ is proportional to the proteome fraction 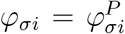 allocated by species *σ* for the uptake and metabolization of resource *i* and *ii)* each resource contributes to the growth of species *σ* through a term 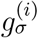 proportional to the uptake rate *J*_*σi*_, so that the total growth rate *g*_*σ*_ of species *σ* can be written as the sum of all the terms 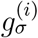. Specifically, we rewrite Eq (1a) as:

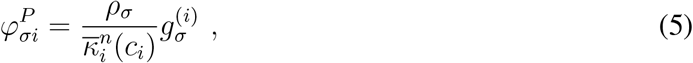

where *ρ* is considered to be species-dependent, 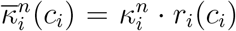 (with *r*_*i*_(*c*_*i*_) = *c*_*i*_/(*K*_*i*_ + *c*_*i*_)), and 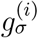 is the contribution to the growth rate of species *σ* due to the uptake of resource *i*, i.e.:

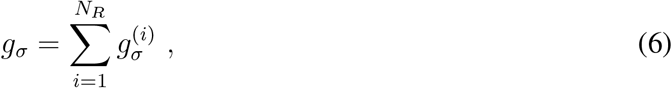

and we generalize Eqs (1a) and (1b) to:

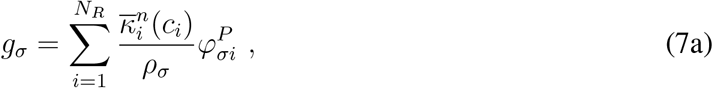

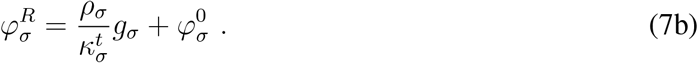

Eq (6) implies that the *N*_*R*_ resources are substitutable (e.g., different carbon sources), otherwise their contribution to the growth rate may satisfy a different equation (e.g., their contributions may be multiplicative rather than additive). We can use Eq (7a) to write Eq (7b) in terms of the fractions 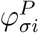. By doing so we get that the normalization condition given by Eq (2) reads:

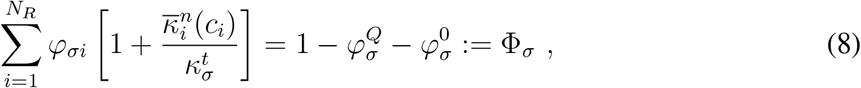

where we have written *φ*_*σi*_ instead of 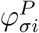 for simplicity and Φ_*σ*_ is the total proteome fraction that species *σ* allocates to metabolism and growth.

We generalize the results of Scott *et al.* to the case of multiple resources by assuming that the uptake rate *J*_*σi*_ of resource *i* per unit biomass is proportional to *φ*_*σi*_, i.e.:

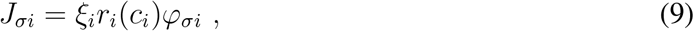

where the proportionality constant *ξ*_*i*_ can be interpreted biologically as the maximum catalytic rate of the enzyme used to metabolize resource *i* (see SI). By comparing Eqs (9) and (5) we can see that the contribution to the growth rate of species *σ* due to the uptake of resource *i* is proportional to its uptake rate, i.e. 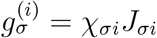with

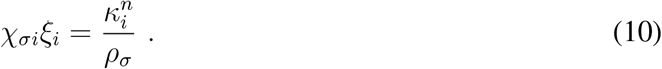

With the considerations above, we obtain the final equations of our model:

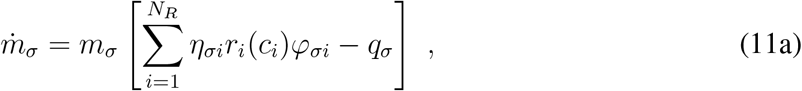

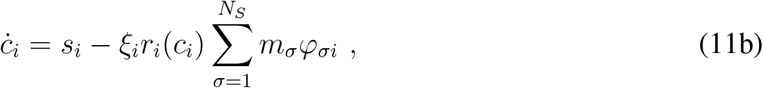

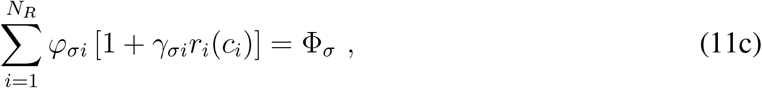

where we have written explicitly 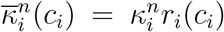 with *r*_*i*_(*c*_*i*_) = *c*_*i*_/(*K*_*i*_ + *c*_*i*_), and we have defined 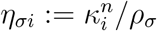 and 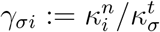 to simplify the notation. Regardless of the particular form of 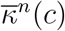 chosen, for our purposes we only need to assume that 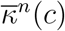 is a monotonically increasing function of *c*, and that 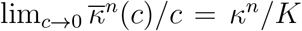 and 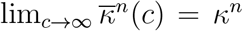. The parameter *ξ*_*i*_ can be interpreted as the maximum catalytic rate of the enzyme used to metabolize resource *i*, and Φ_*σ*_ is the total proteome fraction allocated by species *σ* for nutrient uptake and growth (see Materials and Methods and SI).

The constraint in Eq (11c) is the explicit expression of Eq (2) in our framework, and can be interpreted geometrically: considering species *σ*, the *N*_*R*_-dimensional vector 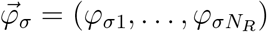 belongs to a hyperplane whose normal vector 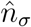 has components 1 + *γ*_*σi*_r_*i*_(*c*_*i*_). This means that as the system evolves, the components of 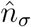 vary with time and therefore the hyperplane to which 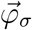 belongs moves in the *N*_*R*_-dimensional space. This is also the reason why the proteome fractions *φ*_*σi*_ must be dynamical variables: the coefficients 1 + *γ*_*σi*_r_*i*_(*c*_*i*_) in Eq (11c) are not fixed, but change with time depending on the system’s dynamics through *r*_*i*_(*c*_*i*_). This implies that for the constraint to be satisfied at all times, the proteome fractions *φ*_*σi*_ *cannot* be fixed but must be, in turn, dynamical variables: an increase (decrease) of 1 + *γ*_*σi*_r_*i*_(*c*_*i*_) must be balanced by a decrease (increase) of some of the *φ*_*σi*_s. This constraint reflects the well known fact that microbes can vary their enzyme synthesis with time and switch between nutrients according to environmental conditions^33,36–38^.

### Dynamics of the proteome fractions *φ*_*σi*_

We call 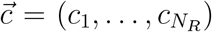 the vector of resource concentrations and define

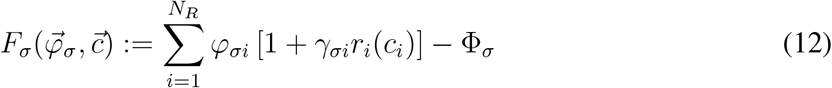

so that the constraint given by Eq (11c) can be written more simply as 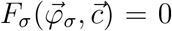. Since this constraint must hold at every instant, any equation for 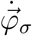 must satisfy

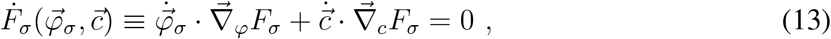

where 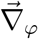 and 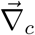 are, respectively, the gradients taken with respect to the components of 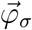 and 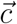. The “minimal” equation for *φ*_*σi*_, i.e. the simplest one (in the sense that it does not introduce extra terms orthogonal to 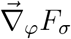, which would lead to a proliferation of new parameters) that satisfies Eq (13), is therefore:

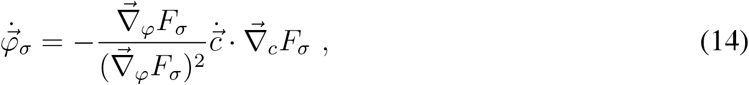

where, however, we are not taking into account the fact that with such an equation some of the *φ*_*σi*_ might become negative with time (see SI for detailed computations on how this can be taken into account).

Microbes are able to switch between nutrients when cultured in mediums containing more than one resource^36^. For this reason, we can implement an adaptive approach^33^ and ask that 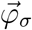 evolves in time so that the growth rate *g*_*σ*_ of species *σ* is maximized respecting the constraint 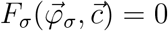, i.e. Eq (11c) is satisfied. In this case the evolution equation for 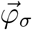 becomes:

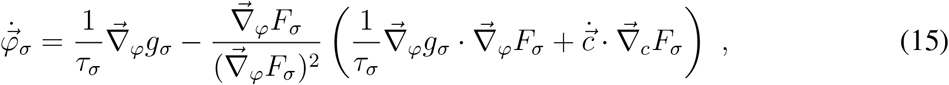

where we have introduced *τ*_*σ*_, the characteristic timescale over which 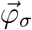 changes^33^ (detailed computations are shown in the SI). We can recover Eq (14) from Eq (15) by sending *τ*_*σ*_ to infinity. Geometrically, Eq (14) represents the case in which 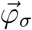 is dragged along by the hyperplane to which it belongs, as the hyperplane moves because of Eq (11c). On the other hand, according to Eq (15) (with small enough values of *τ*_*σ*_) the 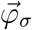 are free to move on the hyperplane to find the maximum instantaneous growth rate compatible with the constraint given by Eq (11c).

In this work we have used a generalization of Eq (15) that ensures *φ*_*σi*_(*t*) ≥ 0 ∀*t*, and varied the values of *τ*_*σ*_ when needed (see SI for details).

### Conditions for coexistence

Evaluating Eqs (11a)–(11c) at stationarity we obtain:

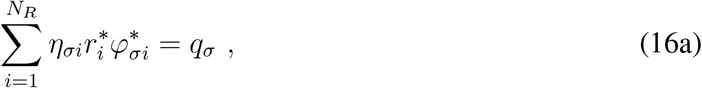

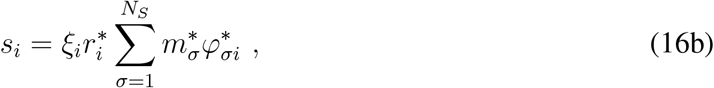

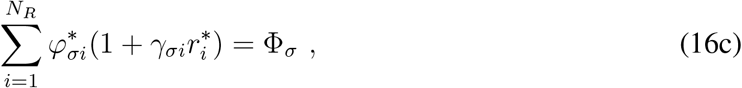

where we are denoting with the symbol “*” the quantities computed at stationarity, and we have assumed 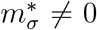. It is easily seen by substitution that a possible solution for 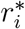 in Eqs (16a) and (16c) is

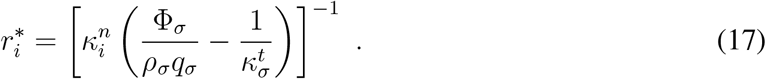

This solution is acceptable only if its right hand side is independent of *σ*, i.e. if

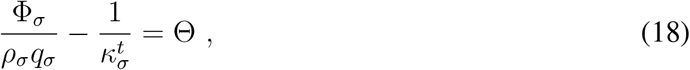

with Θ some given constant independent of *σ*. Using Eqs (17) and (18) in Eq (16c) or (16a) we get

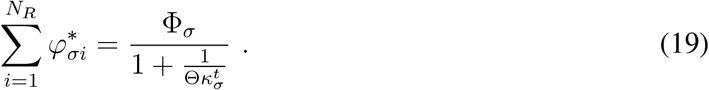

From Eq (17) we have:

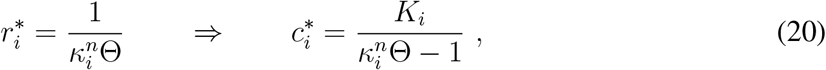

and since we need 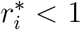 (or equivalently 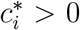), we need 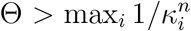. Notice that Eq (18) can be rewritten as

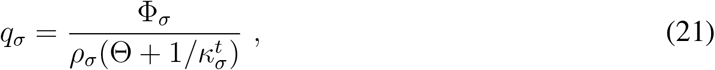

which is the explicit expression of the relationship between *q*_*σ*_ and Φ_*σ*_. Notice that Eq (19) is a consequence of the system’s constraint in Eq (16c), which is Eq (11c) computed at stationarity. Therefore, the expression of the maintenance cost given in Eq (21) is a consequence of the constraint introduced in our consumer-proteome-resource model.

If we now define:

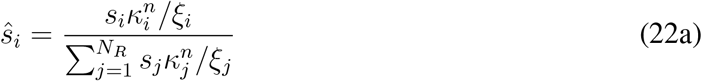

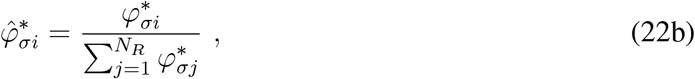

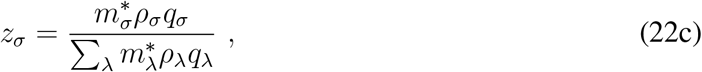

(so that *z*_*σ*_ are positive coefficients that sum to one) Eq (16b) can be indeed rewritten as

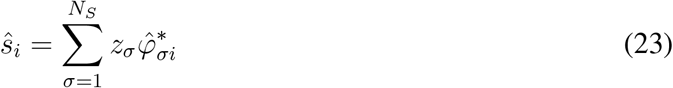

(see SI for the detailed computations).

Since 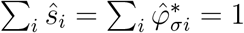, the vectors 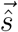and 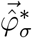 belong to an (*N*_*R*_−1)–dimensional simplex. Furthermore, since *z*_*σ*_ are positive coefficients that sum to one, Eq (23) means that 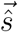 belongs to the convex hull of the vectors 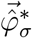. Since Eq (23) derives from requiring that all species have non-null stationary biomasses, we can see how this is the second condition necessary for coexistence. Notice also that Eq (23) depends on the (rescaled) value of *φ*_*σi*_ *at stationarity*, meaning that the proteome fractions *φ*_*σi*_ vary over time as the system evolves to satisfy Eq (23), i.e. to include 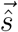 in the convex hull of the vectors 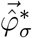.

If we now suppose that *τ*_*σ*_ ≫ 1, so that we can use Eq (14) for the dynamics of *φ*_*σi*_, observing that the *i*-th component of the gradients 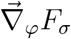 and 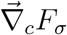 are

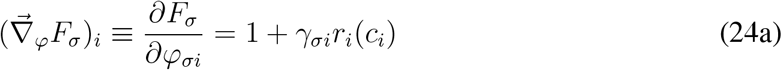

and

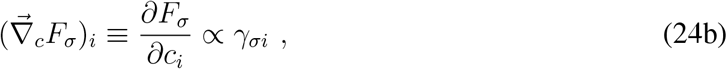

we find that if *γ*_*σi*_ ~ 0 then 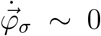 and therefore, if stationarity is reached sufficiently fast, 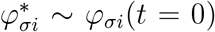. In other words, if the *γ*_*σi*_ are small, the proteome fractions *φ*_*σi*_ at stationarity will be close to their initial values. Therefore in this case, with good approximation, Eq (23) gives the condition for all species to coexist, i.e. 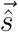 must be inside the convex hull of 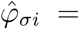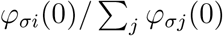. If *γ*_*σi*_ ~ 0 as discussed in the Results section, coexistence will be possible if the components of 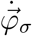 are not too small for a sufficiently long period of time so as to allow them to reach values satisfying Eq (23) and thus for the species to coexist. This can be obtained by using large supply rates *s*_*i*_ so that *r*_*i*_(*c*_*i*_) ~ 1 for a sufficiently long time, as discussed in the Results. Finally, if the ratios *γ*_*σi*_ have larger values the proteome fractions *φ*_*σi*_ will be able to move more quickly.

### Strains used in the experiment

The *Escherichia coli* strains used in our experiment have the same genetic background MG1655. The strains used in the experiments were constructed starting from the ancestor strain 0Y (expressing constitutively the yellow fluorescent protein mVenus from the genome, with genotype attTN7::pRNA1 mVenus) or the ancestor strain 0R (expressing constitutively the red fluorescent protein mKate2Hyb from the genome, with genotype attTN7::pRpsL mKate2Hyb).

Strain 1 was obtained by transforming strain 0Y with the plasmid pR (see Table S.1), which contains the ampicillin resistance cassette, the red fluorescent protein mCherry under the control of the *trc* promoter, a hybrid of the *trp* and *lac* promoters, and the *lac* repressor, *lacI*. The expression of mCherry could thus be induced by adding IPTG, which binds to the repressor encoded by *lacI* allowing the expression of genes promoted by the *trc* promoter (here, mCherry). Because IPTG cannot be metabolized by *E. coli*, its concentration remains constant during our experiment and is unaltered by bacterial growth.

Strain 2 was obtained by transforming strain 0R with the plasmid pAMP (see Table S.1), which was obtained by removing the inducible red fluorescent protein mCherry from plasmid pR using traditional cloning.

Strain 3 was obtained by transforming strain 0R with plasmid pY (see Table S.1), which is identical to plasmid pR, except for the fluorescent protein induced by the *trc* promoter, which is Venus YFP instead of mCherry.

Strain 4 was obtained transforming strain 0Y with plasmid pAMP.

Because all strains had the ampicillin resistance cassette in the plasmids used to transform them, we performed the experiments by adding ampicillin to the medium to prevent contamination and plasmid loss.

### Experimental protocol

The competition assays were performed as follows:

1. The strains were cultured overnight from a stock culture in M63 medium with 1% w/v glucose, and ampicillin. Then, the strains were mixed to perform competition assays aiming for 50:50 relative frequencies.
2. The mixtures were inoculated in a 96-well plate containing M63 medium with 1% w/v glucose and ampicillin at eight different IPTG concentrations: 0*μ*M, 15*μ*M, 30*μ*M, 45*μ*M, 60*μ*M, 75*μ*M, 90*μ*M, 105*μ*M (six technical replicates per concentration).
3. The well plate was covered with a porous rayon film that allowed gas exchange and was cultured for 24 hours at 30°C on a microplate shaker set at 1050rpm.
4. After 24 hours, the plate was reinoculated in a new 96-well plate with fresh medium (with the appropriate concentrations of IPTG in each well) with a dilution factor of 100. The new plate was cultured for another cycle at 30°C for 24 hours with constant shaking at 1050rpm, while the old one was diluted with a dilution factor of 2000 to be analyzed at the flow cytometer.

### IPTG calibration and computation of the normalized protein production rate

We measured how the fluorescence intensity of individual cells, a proxy for the total amount of fluorescent protein produced, varied as a function of the IPTG concentration. To do so, we inoculated strains 1 and 3 in a 96-well plate containing M63 minimal medium with ampicillin, 1% w/v glucose and the same IPTG concentrations used in our experimental protocol (six technical replicates per concentration, per strain). The plate was incubated at 30°C for 8 hours with constant shaking at 1050rpm. At times *t* = 4h and *t* = 8h after inoculation we measured at the flow-cytometer the mean fluorescence intensity of cells due to the induced fluorescent proteins at the various concentrations of IPTG (Figures 3d and 3e). From these data, we estimated the normalized fluorescent protein production rate as follows.

We call *k*(*C*_*I*_) the rate at which the fluorescence of the inducible protein increases when cells are exposed to a concentration *C*_*I*_ of IPTG, and we call *d*_*FP*_ the fluorescent protein degradation rate. The fluorescent intensity 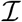 of a cell (due to the production of the IPTG-inducible fluorescent protein) in between two successive cell divisions thus satisfies 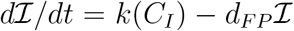. At a cell division event, the fluorescent intensity of a cell is reduced by a factor 2. Indicating with 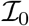 the cell’s fluorescent intensity at the first measurement time (*t* = 4h), it can be shown (see SI) that according to this model the cell’s fluorescent intensity changes with time as:

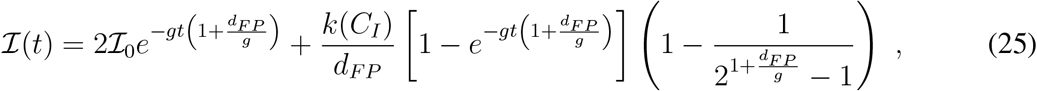

where *g* is the cell’s growth rate. Fluorescent proteins have small degradation rates compared to the cellular growth rate, so assuming *d*_*FP*_ ≪ *g* we can approximate Eq 25 as:

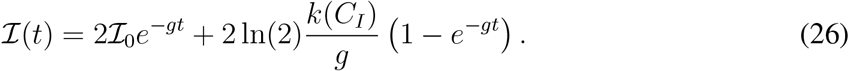

We used Eq 26 and the data Figures 3d and 3e to compute the quantity *k*(*C*_*I*_). Because the absolute value of *k*(*C*_*I*_) depends on the arbitrary units returned by the flow cytometer (the intensity 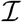 is measured as a cell’s pulse area at the flow cytometer), we normalized the values of *k*(*C*_*I*_) dividing them by the mean fluorescent intensity 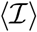 of cells measured in the absence of IPTG at the first measurement in the calibration experiment (see Figures 3d and 3e). Such a normalization affects only the absolute value of such rates, and not their relative magnitude. This also means that the normalized production rates shown for the two experiments in Figure 3 cannot be compared directly. The normalized 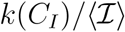 are the protein production rates (with dimensions 1/time) reported in Figure 3.

### Estimation of the selection coefficient 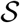

Applying Eqs (11a)–(11c) to the case of two populations and one resource, we obtain:

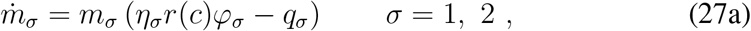

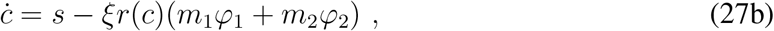

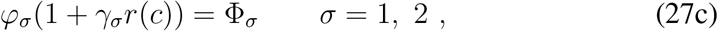

where now Eq (27c) gives the explicit expression of the (only) proteome fraction *φ*_*σ*_ as a function of the resource concentration. Because the ancestors of our two strains (i.e., strains 1 and 2 without the plasmids conferring ampicillin resistant and the inducible RFP, see Materials and Methods) have equal growth rates (see Figures S.8 and S.9), we set *η*_1_ = *η*_2_ = *η*, *q*_1_ = *q*_2_ = *q* and *γ*_1_ = *γ*_2_ = *γ* in Eqs (27a)–(27c). Note that, instead, Φ_1_ ≠ Φ_2_ because the proteome allocation of strain 1 could be varied experimentally and because the plasmids introduced in the ancestor strains have different maintenance costs. Given that cells in the experiment are grown in nutrient-rich conditions, we assume that the maintenance cost is negligible, i.e. *q* ⋍ 0. Furthermore, because most of the dynamics (i.e., the relative change in abundance of the two strains) occurs in the early phases of growth when glucose is abundant, we assume that *r*(*c*) ≈ 1 at all times so that we can neglect Eq (27b) and we are left with:

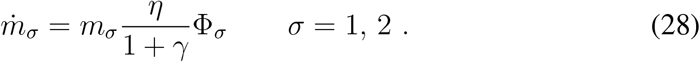

Notice again that this expression, and in particular the fact that the growth rate of species *σ* is proportional to Φ_*σ*_, is a consequence of the constraint in Eq (27c).

We therefore have that the expression of the selective advantage 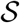 is:

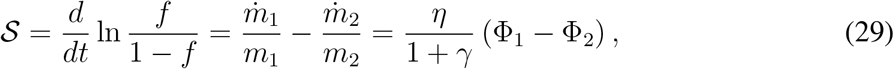

where *f* = *m*_1_/(*m*_1_ + *m*_2_) and 1 − *f* = *m*_2_/(*m*_1_ + *m*_2_) are the relative abundances (or “frequencies”) of strain 1 and strain 2, respectively. In the SI we show that if we lift the assumption that *r*(*c*) = 1 at all times, we obtain:

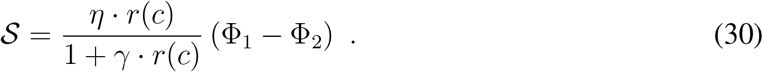

### Evaluation of the ratios Φ_1_/Φ_2_ and Φ_3_/Φ_4_

Consider the competition assay with strains 1 and 2 (the results are the same also for the competition assay between strains 3 and 4, after all subscripts are appropriately changed). For a given IPTG concentration *C*_*I*_, from Eq (28) the growth rate of strain 1 is *g*_1_[*k*(*C*_*I*_)] = Φ_1_[*k*(*C*_*I*_)] · *r*(*c*)*η*/(1 + *γr*(*c*)) (where we have inserted explicitly the dependence on *r*(*c*), and *k*(*C*_*I*_) is the protein production rate induced by *C*_*I*_). Dividing 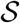 in Eq (30) by *g*_1_, for any value of *r*(*c*) we obtain:

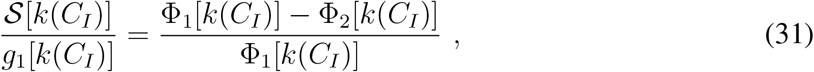

which is easily rearranged into:

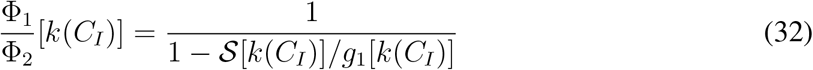

(which are the values plotted in Figure 3c). Notice that, this result does *not* depend on the assumption that *r*(*c*) = 1 at all times, i.e. Eq (32) is valid for any value of *r*(*c*).

### Selective advantage and proteome fraction allocated to the useless RFP

Because we did not measure RNA/protein ratios in our experiments, we can only estimate the values of *κ*^*t*^ and *κ*^*n*^ by taking them from the literature for *E. coli* strains grown at 30°C in conditions similar to our experiments. Rosset *et al.*^39, 40^ measured the RNA/protein ratio of several *E. coli* strains grown at 30°C in M63 medium. Using their data and the relationship^22^ *r* = *r*_0_ + *g/κ^t^*, where *r* is the RNA/protein ratio and *r*_0_ a constant, we can estimate the translational capacity as *κ*^*t*^ = 3.0 ± 0.5 *μ*g protein/*μ*g RNA · 1/h (mean±SD). An estimate for the nutritional capacity *κ*^*n*^, instead, can be obtained via the equation^22^ *g* = *g*_max_*κ^n^/*(*κ*^*n*^ + *κ*^*t*^), where *g*_max_ is the maximum growth rate obtainable by our strain at a given temperature (for us, 30°C), when nutrients are abundant. Van Derlinden and Van Impe^41^ report a maximum growth rate *g*_max_ ≈ 1.2 1/h for *E. coli* MG1655 grown at 30°C in rich medium with glucose (no error estimate was reported). Solving for *κ*^*n*^ and using the growth rate value *g* measured for strain 1 in the absence of IPTG, we find *κ*^*n*^ = 1.2 ± 0.2 *μ*g protein/*μ*g RNA · 1/h. These values allow us to estimate *γ* = *κ*^*n*^/*κ*^*t*^ = 0.4 ± 0.1 and *η* = *κ*^*n*^/*ρ* = 1.57 ± 0.07 1/h using the value for *ρ* = 0.76 *μ*g protein/*μ*g RNA · 1/h reported in Scott *et al.*^22^. With these estimations, from the expression of the selective advantage in Eq (30) we have that a 1% difference in proteome allocation for metabolism and growth between the two strains (i.e., Φ_1_ − Φ_2_ = 1%) leads to 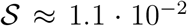. Finally, with these calculations we can estimate the maximum percentage of proteome max *φ*_*iRFP*_ and max *φ*_*iY FP*_ allocated to the production of, respectively, the inducible red and yellow proteins in our two experiments. In particular, for the first experiment we have max 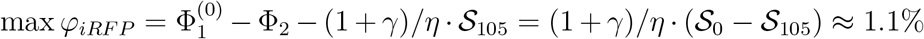 (where 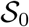 and 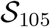 are, respectively, the mean selection coefficients in the 0 *μ*M and 105 *μ*M IPTG treatments). For the experiment involving strains 3 and 4, using the same procedure we find max *φ*_*iY FP*_ ≈ 0.4%. Of course, given that we had to rely on measurements taken from the literature, these should be regarded as only rough estimates.

## Acknowledgements

We thank David R Nelson and Andrew W Murray for hosting L. P.-M. during the experiment and the initial development of the model and for insightful comments and suggestions. We thank Daniel Eaton for providing the ancestor bacterial strains used in the experiment. A. M. and L. P.-M. acknowledge the Cariparo Foundation for funding. S. S. acknowledges the University of Padua for STARS ReACT grant. A. G. was supported by research fellowships from the Swiss National Science Foundation, Projects P2ELP2_168498, P400PB_180823 and P400PB_180823 / 2.

