## Supplementary Information for "Constrained proteome allocation affects coexistence in models of competitive microbial communities"

#### CONTENTS

I. The consumer-proteome-resource model 2

A. Biological interpretation of $\xi_i$ in Eq (11b) 2

B. Dynamics of $\varphi_{\sigma i}$ – complete expression of Eq (15) 2

C. Conditions for coexistence – derivation of Eq (23) 2

D. Parameters used in the simulations 3

II. Experiment 3

A. Computation of the normalized protein production rate 3

B. Data 4

1. Time series 4

2. OD curves and growth rates 4

C. A more detailed model 4

References 5

Figures and Tables 6

### I. THE CONSUMER-PROTEOME-RESOURCE MODEL

#### A. Biological interpretation of $\xi_i$ in Eq (11b)

Scott et al. [1, Supporting Online Material] give a microscopic interpretation of the nutritional capacity $\kappa_i^n$ by stating that the growth rate of a microbial species is given by $g = qJ$, where $J$ is the uptake rate of the (only) resource per unit biomass, and $q$ is a proportionality constant that depends on the properties of the nutrient (e.g., how much energy its metabolization can generate). They then assume that there is only one bottleneck enzyme $E$ for the growth of the microbial species, and write:

$$J = k_E r(c) \varphi_E \ , \tag{S.1}$$

where $k_E$ is the maximal catalytic rate of enzyme $E$, $r(c)$ is Monod's function, and $\varphi_E$ is the fraction of the proteome occupied by the enzyme $E$. Compared to our formalism in the general case of $N_S$ species and $N_R$ resources (so that $J \to J_{\sigma i}$ and $g \to g_\sigma$), we can identify $r(c) \to r_i(c_i)$, $\varphi_E \to \varphi_{\sigma i}$, $q \to \chi_{\sigma i}$ and finally $k_E \to \xi_i$.

#### B. Dynamics of $\varphi_{\sigma i}$ – complete expression of Eq (15)

In components, Eq (15) is written as follows:

$$\dot{\varphi}_{\sigma i} = \frac{1}{\tau_\sigma}\frac{\partial g_\sigma}{\partial \varphi_{\sigma i}} - \frac{\partial F_\sigma/\partial \varphi_{\sigma i}}{\sum_{k=1}^{N_R} \left(\partial F_\sigma/\partial \varphi_{\sigma k}\right)^2} \sum_{j=1}^{N_R} \left( \frac{1}{\tau_\sigma}\frac{\partial g_\sigma}{\partial \varphi_{\sigma j}}\frac{\partial F_\sigma}{\partial \varphi_{\sigma j}} + \dot{c}_j \frac{\partial F_\sigma}{\partial c_j} \right) \ . \tag{S.2}$$

To make sure that $\varphi_{\sigma i}$ doesn't become negative, we use the following approach. We rewrite Eq (S.2) in terms of an auxiliary variable $\psi_{\sigma i}$ instead of $\varphi_{\sigma i}$:

$$\dot{\psi}_{\sigma i} = \frac{1}{\tau_\sigma}\frac{\partial g_\sigma}{\partial \psi_{\sigma i}} - \frac{\partial F_\sigma/\partial \psi_{\sigma i}}{\sum_{k=1}^{N_R} \left(\partial F_\sigma/\partial \psi_{\sigma k}\right)^2} \sum_{j=1}^{N_R} \left( \frac{1}{\tau_\sigma}\frac{\partial g_\sigma}{\partial \psi_{\sigma j}}\frac{\partial F_\sigma}{\partial \psi_{\sigma j}} + \dot{c}_j \frac{\partial F_\sigma}{\partial c_j} \right) \ , \tag{S.3}$$

and then we define $\varphi_{\sigma i} = \mathcal{F}(\psi_{\sigma i})$, where $\mathcal{F}$ is a positive function, i.e. $\mathcal{F}(x) > 0 \ \forall x$. This way, Eq (S.3) becomes:

$$\dot{\varphi}_{\sigma i} = \mathcal{F}'(\psi_{\sigma i})^2 \left[ \frac{1}{\tau_\sigma}\frac{\partial g_\sigma}{\partial \varphi_{\sigma i}} - \frac{\partial F_\sigma/\partial \varphi_{\sigma i}}{\sum_{k=1}^{N_R} \left(\partial F_\sigma/\partial \varphi_{\sigma k}\right)^2 \left(\mathcal{F}'(\psi_{\sigma k})\right)^2} \sum_{j=1}^{N_R} \left( \frac{\mathcal{F}'(\psi_{\sigma j})^2}{\tau_\sigma}\frac{\partial g_\sigma}{\partial \varphi_{\sigma j}}\frac{\partial F_\sigma}{\partial \varphi_{\sigma j}} + \dot{c}_j \frac{\partial F_\sigma}{\partial c_j} \right) \right] \ . \tag{S.4}$$

If we choose $\mathcal{F}(x) = x^2/4$ (any other choice leads to qualitatively identical results), so that $\mathcal{F}'(\psi_{\sigma i})^2 = \varphi_{\sigma i}$, and taking into account that:

$$\frac{\partial g_\sigma}{\partial \varphi_{\sigma i}} = \eta_{\sigma i} r_i(c_i) \ , \qquad \frac{\partial F_\sigma}{\partial \varphi_{\sigma i}} = 1 + \gamma_{\sigma i} r_i(c_i) \ , \qquad \frac{\partial F_\sigma}{\partial c_i} = \varphi_{\sigma i}\gamma_{\sigma i}\frac{K_i}{(c_i + K_i)^2} \ , \tag{S.5}$$

we have that the final expressions of the equation for $\dot{\varphi}_{\sigma i}$ (i.e., Eq (15)), in components, is:

$$\dot{\varphi}_{\sigma i} = \varphi_{\sigma i} \left[ \frac{\eta_{\sigma i} r_i(c_i)}{\tau_\sigma} - \frac{(1 + \gamma_{\sigma i} r_i(c_i))}{\sum_{k=1}^{N_R} \varphi_{\sigma k}(1 + \gamma_{\sigma k} r_k(c_k))^2} \sum_{j=1}^{N_R} \varphi_{\sigma j} \left( \frac{\eta_{\sigma j} r_j(c_j)}{\tau_\sigma}(1 + \gamma_{\sigma j} r_j(c_j)) + \gamma_{\sigma j}\frac{K_j}{(c_j + K_j)^2}\dot{c}_j \right) \right] \ . \tag{S.6}$$

#### C. Conditions for coexistence – derivation of Eq (23)

Plugging Eq (17) into Eq (16b) we obtain:

$$s_i \frac{\kappa_i^n}{\xi_i} = \frac{1}{\Theta}\sum_{\sigma=1}^{N_S} m_\sigma^* \varphi_{\sigma i}^* \ , \tag{S.7}$$

and by summing this equation over $i$ on both sides, we get

$$\sum_{i=1}^{N_R} s_i \frac{\kappa_i^n}{\xi_i} = \sum_{\sigma=1}^{N_S} m_\sigma^* \frac{1}{\Theta} \sum_{i=1}^{N_R} \varphi_{\sigma i}^* = \sum_{\sigma=1}^{N_S} m_\sigma^* \rho_\sigma q_\sigma \ . \tag{S.8}$$

If we now define

$$\hat{s}_i := \frac{s_i \kappa_i^n/\xi_i}{\sum_{j=1}^{N_R} s_j \kappa_j^n/\xi_j} \ , \qquad\qquad \hat{\varphi}^*_{\sigma i} := \frac{\varphi^*_{\sigma i}}{\sum_{j=1}^{N_R} \varphi^*_{\sigma j}} \tag{S.9}$$

(so that $\sum_i \hat{s}_i = \sum_i \hat{\varphi}_{\sigma i} = 1$), Eq (S.7) becomes:

$$\hat{s}_i = \sum_{\sigma=1}^{N_S} \frac{m^*_\sigma \rho_\sigma q_\sigma}{\sum_{\lambda=1}^{N_S} m^*_\lambda \rho_\lambda q_\lambda} \hat{\varphi}^*_{\sigma i} \ , \tag{S.10}$$

that is exactly Eq (23) of the Main Text once we define

$$z_\sigma := \frac{m^*_\sigma \rho_\sigma q_\sigma}{\sum_{\lambda=1}^{N_S} m^*_\lambda \rho_\lambda q_\lambda} \ , \tag{S.11}$$

which are indeed positive coefficients such that $\sum_{\sigma=1}^{N_S} z_\sigma = 1$.

##### D. Parameters used in the simulations

**Figure 5:** The parameters used to run the simulations shown in Figure 5 of the Main Text were drawn randomly from the following distributions: $m_\sigma(0) \in \mathcal{U}[1,5]$ $\mu$g of biomass/mL (which is in the order of $\approx 10^6$ cells/mL, assuming an average mass of $\approx 10^{-12}$ g for an *E. coli* cell), with $\mathcal{U}$ the uniform distribution, $c_i \in \mathcal{U}[0.5 \cdot 10^4, 1.5 \cdot 10^4]$ $\mu$g of resource/mL (which is in the order of a concentration of roughly 1% w/v), $\kappa_i^n \in \mathcal{U}[10,50]$ $\mu$g protein/$\mu$g RNA $\cdot$ 1/h, $\kappa_\sigma^t \in \mathcal{U}[1,5]$ $\mu$g protein/$\mu$g RNA $\cdot$ 1/h (so that $\gamma_{\sigma i} \sim O(0.1)$), $\rho_\sigma \in \mathcal{U}[0.5, 0.9]$ $\mu$g protein/$\mu$g RNA, $s_i \in \mathcal{U}[0.5 \cdot 10^4, 2 \cdot 10^4]$ $\mu$g of resource/mL $\cdot$ 1/h, $\xi_i \in \mathcal{U}[10^3, 1.5 \cdot 10^3]$ 1/h (which corresponds to $\approx 0.35$ 1/s, the catalytic rate of the slowest enzyme involved in the glycolytic pathway of *E. coli*, as reported by Gameiro et al. [2]), $K_i \in \mathcal{U}[5 \cdot 10^2, 10^3]$ $\mu$g/mL (which is in the order of $\approx 8 \cdot 10^2$ $\mu$g/L as reported by Seen et al. [3] for *E. coli* at 30°C), $\tau_\sigma \in \mathcal{U}[10^4, 5 \cdot 10^4]$ 1/h (so that $\tau_\sigma \gg 1$), $\Phi_\sigma \in \mathcal{U}[0.45, 0.55]$, $\varphi_{\sigma i}$ were drawn randomly so that they satisfy $\sum_i \varphi_{\sigma i}(1 + \gamma_{\sigma i} r_i(c_i(0))) = \Phi_\sigma$, and $\Theta = 5$.

**Figure 6:** The parameters used to run the simulations shown in Figure 6 of the Main Text were drawn randomly from the following distributions: $m_\sigma(0) \in \mathcal{U}[1,5]$ $\mu$g of biomass/mL, $c_i \in \mathcal{U}[0.5 \cdot 10^4, 1.5 \cdot 10^4]$ $\mu$g of resource/mL, $\kappa_i^n \in \mathcal{U}[5,10]$ $\mu$g protein/$\mu$g RNA $\cdot$ 1/h, $\kappa_\sigma^t \in \mathcal{U}[1,5]$ $\mu$g protein/$\mu$g RNA $\cdot$ 1/h (so that $\gamma_{\sigma i} \sim O(2)$), $\rho_\sigma \in \mathcal{U}[0.6, 0.8]$ $\mu$g protein/$\mu$g RNA, $s_i \in \mathcal{U}[0.5 \cdot 10^4, 10^4]$ $\mu$g of resource/mL $\cdot$ 1/h, $\xi_i \in \mathcal{U}[10^3, 1.5 \cdot 10^3]$ 1/h, $K_i \in \mathcal{U}[5 \cdot 10^2, 10^3]$ $\mu$g/mL, $\tau_\sigma \in \mathcal{U}[10^4, 5 \cdot 10^4]$ 1/h (so that $\tau_\sigma \gg 1$), $\Phi_\sigma \in \mathcal{U}[0.45, 0.55]$, $\varphi_{\sigma i}$ were drawn randomly so that they satisfy $\sum_i \varphi_{\sigma i}(1 + \gamma_{\sigma i} r_i(c_i(0))) = \Phi_\sigma$, and $\Theta = 5$.

**Figure 7:** The parameters used to run the simulations shown in Figure 7 of the Main Text were drawn randomly from the following distributions: $m_\sigma(0) \in \mathcal{U}[1,5]$ $\mu$g of biomass/mL, $c_i \in \mathcal{U}[0.5 \cdot 10^4, 1.5 \cdot 10^4]$ $\mu$g of resource/mL, $\kappa_i^n \in \mathcal{U}[2,5]$ $\mu$g protein/$\mu$g RNA $\cdot$ 1/h, $\kappa_\sigma^t \in \mathcal{U}[1,4]$ $\mu$g protein/$\mu$g RNA $\cdot$ 1/h (so that $\gamma_{\sigma i} \sim O(1)$), $\rho_\sigma \in \mathcal{U}[0.6, 0.8]$ $\mu$g protein/$\mu$g RNA, $s_i \in \mathcal{U}[0.5 \cdot 10^4, 2 \cdot 10^4]$ $\mu$g of resource/mL $\cdot$ 1/h, $\xi_i \in \mathcal{U}[10^3, 1.5 \cdot 10^3]$ 1/h, $K_i \in \mathcal{U}[5 \cdot 10^2, 10^3]$ $\mu$g/mL, $\tau_\sigma \in \mathcal{U}[1,5]$ 1/h, $\Phi_\sigma \in \mathcal{U}[0.45, 0.55]$, $\varphi_{\sigma i}$ were drawn randomly so that they satisfy $\sum_i \varphi_{\sigma i}(1 + \gamma_{\sigma i} r_i(c_i(0))) = \Phi_\sigma$, and $\Theta = 5$.

#### II. EXPERIMENT

##### A. Computation of the normalized protein production rate

Let us call $k(C_I)$ the production rate of the induced fluorescent protein when cells are exposed to an IPTG concentration of $C_I$, and let us call $d_{FP}$ the protein degradation rate. If we call $\mathcal{I}$ the fluorescent intensity of a cell due to the induced protein, between two consecutive cell divisions $\mathcal{I}$ will satisfy:

$$\frac{d\mathcal{I}}{dt} = k(C_I) - d_{FP}\mathcal{I} \ , \tag{S.12}$$

whose solution is:

$$\mathcal{I}(t) = \mathcal{I}_0 e^{-d_{FP}t} + \frac{k(C_I)}{d_{FP}} \left(1 - e^{-d_{FP}t}\right) \ . \tag{S.13}$$

If we now call $\tau$ the doubling time and we take into account that the $\mathcal{I}$ is halved after every cellular division, the fluorescent intensity $\mathcal{I}_i$ at the $i$-th division will be:

$$\mathcal{I}_1 = \mathcal{I}_0 e^{-d_{FP}\tau} + \frac{k(C_I)}{d_{FP}}(1 - e^{-d_{FP}\tau}) \tag{S.14a}$$

$$\mathcal{I}_2 = \frac{\mathcal{I}_1}{2} e^{-d_{FP}\tau} + \frac{k(C_I)}{d_{FP}}(1 - e^{-d_{FP}\tau}) = \frac{\mathcal{I}_0}{2} e^{-d_{FP}2\tau} + \frac{k(C_I)}{d_{FP}}(1 - e^{-d_{FP}\tau})\left(1 + \frac{1}{2}e^{-d_{FP}\tau}\right) \tag{S.14b}$$

$$\mathcal{I}_3 = \frac{\mathcal{I}_2}{2} e^{-d_{FP}\tau} + \frac{k(C_I)}{d_{FP}}(1 - e^{-d_{FP}\tau}) = \frac{\mathcal{I}_0}{4} e^{-d_{FP}3\tau} + \frac{k(C_I)}{d_{FP}}(1 - e^{-d_{FP}\tau})\left(1 + \frac{1}{2}e^{-d_{FP}\tau} + \frac{1}{4}e^{-2d_{FP}\tau}\right) \tag{S.14c}$$

$$\vdots$$

$$\mathcal{I}_i = \frac{\mathcal{I}_0}{2^{i-1}} e^{-d_{FP}\tau i} + \frac{k(C_I)}{d_{FP}}(1 - e^{-d_{FP}\tau}) \sum_{\ell=1}^{i} \frac{e^{-(\ell-1)d_{FP}\tau}}{2^{\ell-1}} \ . \tag{S.14d}$$

If we now define $\tau i = t$, and take into account that $\tau = \ln 2/g$ (with $g$ the cell's growth rate), and we use the fact that

$$\sum_{\ell=1}^{i} \frac{e^{-(\ell-1)d_{FP}\tau}}{2^{\ell-1}} = \frac{2e^{d_{FP}\tau}(1 - 2^{-i}e^{-d\tau i})}{2e^{d_{FP}\tau} - 1} \ , \tag{S.15}$$

and we can rewrite Eq (S.14d) as:

$$\mathcal{I}(t) = 2\mathcal{I}_0 e^{-gt\left(1+\frac{d_{FP}}{g}\right)} + \frac{k(C_I)}{d_{FP}}\left[1 - e^{-gt\left(1+\frac{d_{FP}}{g}\right)}\right]\left(1 - \frac{1}{2^{1+\frac{d_{FP}}{g}} - 1}\right) \tag{S.16}$$

which is Eq (25) in the Main Text.

#### B. Data

##### 1. *Time series*

Figures S.2 and S.3 show the time series of $\ln f/(1-f)$ for both the experiments shown in Figure 3 of the Main Text. We can see that the trend of the first day is in contrast with the rest of the experiment (i.e., $\ln f/(1-f)$ decreases in the first 24h and then increases, or vice versa). We found out that this is due to the fact at 0h some flow-cytometry data are very close to the experimental noise, and therefore the two strains are difficult to resolve. The selective advantages have therefore been computed by fitting each curve with a linear function between day 1 and day 6 (i.e., between 24h and 144h).

##### 2. *OD curves and growth rates*

Figures S.4 and S.6 show the growth curves of strains 1 and 3 for the different IPTG concentrations. Figures S.5 and S.7, then, show the values of the growth rates obtained from these curves.

#### C. A more detailed model

When applying the model to the data, we have assumed that $r(c) = 1$ at all times to obtain the expression of the selective advantage in Eq (29). However, in our experiment this was not the case since the density of the cells saturated well before the following re-inoculation in fresh medium was made (i.e., 24 hours).[1] Here we show that by taking into account the fact that cells can reach saturation before the following re-inoculation, our results remain unchanged.

A model better suited to describe the experiments would be as follows. The temporal dynamics of biomass and glucose concentration between two consecutive dilutions satisfies:

$$\dot{m}_\sigma = m_\sigma \eta_\sigma r(c) \varphi_\sigma \qquad \sigma = 1, 2 \tag{S.17a}$$

$$\dot{c} = -r(c)(m_1 \eta_1 \varphi_1 + m_2 \eta_2 \varphi_2) \ , \tag{S.17b}$$

where $c(t)$ is the concentration of glucose at time $t$ and $r(c) = c/(c+K)$ is Monod's function. This model is somewhat similar to MacArthur's consumer-resource model, with the difference that there is no mortality term in Eq (S.17a): between two consecutive dilutions, the biomass $m_\sigma$ of strain $\sigma$ will grow as long as there is glucose available, and because $r(0) = 0$ the strains will stop growing (i.e. they will enter the stationary phase) once glucose runs out.

We now make the following approximation: we assume that, after every reinoculation, glucose is initially abundant (i.e. $r \sim 1$) and that the transition of $r(c)$ from 1 to 0 as $c$ decreases is abrupt, which happens if $K$ is sufficiently small. In other words, we assume that $K$ is sufficiently small so that $r(c) \sim 1$ until a given time $T$ (the instant at which glucose is completely depleted), when $r(c)$ abruptly goes to zero (i.e., $r(c(t)) \sim \Theta(T-t)$ with $\Theta$ the Heaviside's step function). This means that after a reinoculation $m_\sigma$ will grow exponentially for a time interval of length $T$, after which it will stop until the next dilution. If we still set $\eta_1 = \eta_2 = \eta$ and $\gamma_1 = \gamma_2 = \gamma$ and call $D$ the dilution factor between reinoculations, we have that the biomass $m_\sigma^{(N)}$ of strain $\sigma$ at the $N$-th dilution is ($m_\sigma^{(0)}$ being the biomass at the initial inoculation):

$$m_\sigma^{(1)} = m_\sigma^{(0)} \exp\left(\frac{\eta}{1+\gamma}\Phi_\sigma T\right) \tag{S.18a}$$

$$m_\sigma^{(2)} = \frac{m_\sigma^{(1)}}{D} \exp\left(\frac{\eta}{1+\gamma}\Phi_\sigma T\right) = \frac{m_\sigma^{(0)}}{D} \exp\left(2\frac{\eta}{1+\gamma}\Phi_\sigma T\right) \tag{S.18b}$$

$$m_\sigma^{(3)} = \frac{m_\sigma^{(2)}}{D} \exp\left(\frac{\eta}{1+\gamma}\Phi_\sigma T\right) = \frac{m_\sigma^{(0)}}{D} \exp\left(3\frac{\eta}{1+\gamma}\Phi_\sigma T\right) \tag{S.18c}$$

$$\vdots$$

$$m_\sigma^{(N)} = \frac{m_\sigma^{(0)}}{D^{N-1}} \exp\left(N\frac{\eta}{1+\gamma}\Phi_\sigma T\right) \ . \tag{S.18d}$$

Computations analogous to those shown in the Methods section lead to:

$$\ln\frac{f^{(N)}}{(1-f)^{(N)}} = \ln\frac{m_1^{(0)}}{m_2^{(0)}} + NT\frac{\eta}{1+\gamma}(\Phi_1 - \Phi_2) \ , \tag{S.19}$$

which gives the same expression for the selection coefficients after deriving with respect to $NT$.

[1] M. Scott, C. W. Gunderson, E. M. Mateescu, Z. Zhang, and T. Hwa, Interdependence of cell growth and gene expression, Science **330**, 10.1177/42.6.8189037 (2010).

[2] D. Gameiro, M. Pérez-Pérez, G. Pérez-Rodríguez, G. Monteiro, N. F. Azevedo, and A. Lourenço, Computational resources and strategies to construct single-molecule metabolic models of microbial cells, Briefings in Bioinformatics **17**, 863 (2015).

[3] H. Senn, U. Lendenmann, M. Snozzi, G. Hamer, and T. Egli, The growth of escherichia coli in glucose-limited chemostat cultures: a re-examination of the kinetics, Biochimica et Biophysica Acta (BBA) - General Subjects **1201**, 424 (1994).

[1] The typical growth rate of the strains, estimated from growth curves measured in the same experimental conditions used for the competition assays, is 0.3 1/h. The competition assays started from a cellular density of $\sim 8 \cdot 10^6$ cells/mL, thus if the growth was exponential the density after 24 hours would have been be $\sim 1.4 \cdot 10^{10}$, which is much higher than the typical density ($\sim 10^9$ cells/mL) that *E. coli* cells reach at saturation. With a growth rate of 0.3 1/h, the time needed to reach a cellular density that is hundredfold the initial one (and therefore the time needed to reach saturation after a re-inoculation) is approximately 15.4 h.

| Strain or plasmid name | Genotype/description | Name in collection |
|---|---|---|
| Plasmid pR | pTrc99A with mCherry introduced in the multiple cloning site. $Amp^R$, $lacI^Q$, pBR322 origin, pTrc_mCherry | pbAG3 |
| Plasmid pY | pTrc99A with Venus YFP introduced in the multiple cloning site. $Amp^R$, $lacI^Q$, pBR322 origin, pTrc_Venus YFP | pbAG2 |
| Plasmid pAMP | Largest digestion of pR with restriction enzyme SphI, ligated with ligase. $Amp^R$, pBR322 origin | pbAG9 |
| Strain 0R | Markerless, chromosomally integrated insertion attTN7::pRpsL_mKate2Hyb | bAG11 |
| Strain 0Y | Markerless, chromosomally integrated insertion attTN7::pRNA1_mVenus | bAG13 |
| Strain 1 | 0Y transformed with pR | bAG17 |
| Strain 2 | 0R transformed with pAMP | bAG25 |
| Strain 3 | 0R transformed with pY | bAG16 |
| Strain 4 | 0Y transformed with pAMP | bAG24 |

**TABLE S.1:** Information on the strains and plasmids used in our experiments

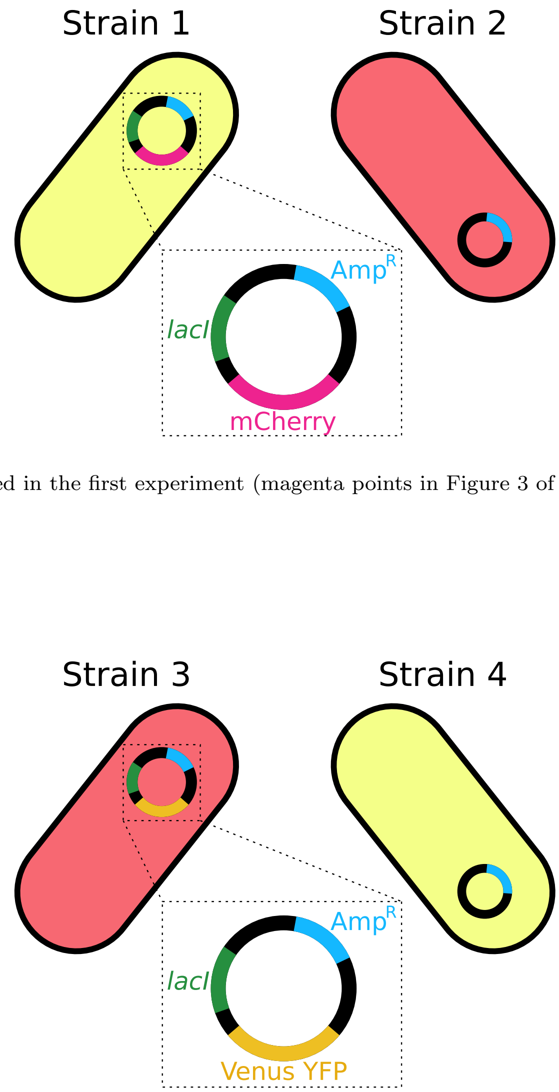


**(a)** Strains used in the first experiment (magenta points in Figure 3 of the Main Text).



**(b)** Strains used in the second experiment (cyan points in Figure 3 of the Main Text).

**FIG. S.1:** Schematic representation of the strains used in both our experiments.

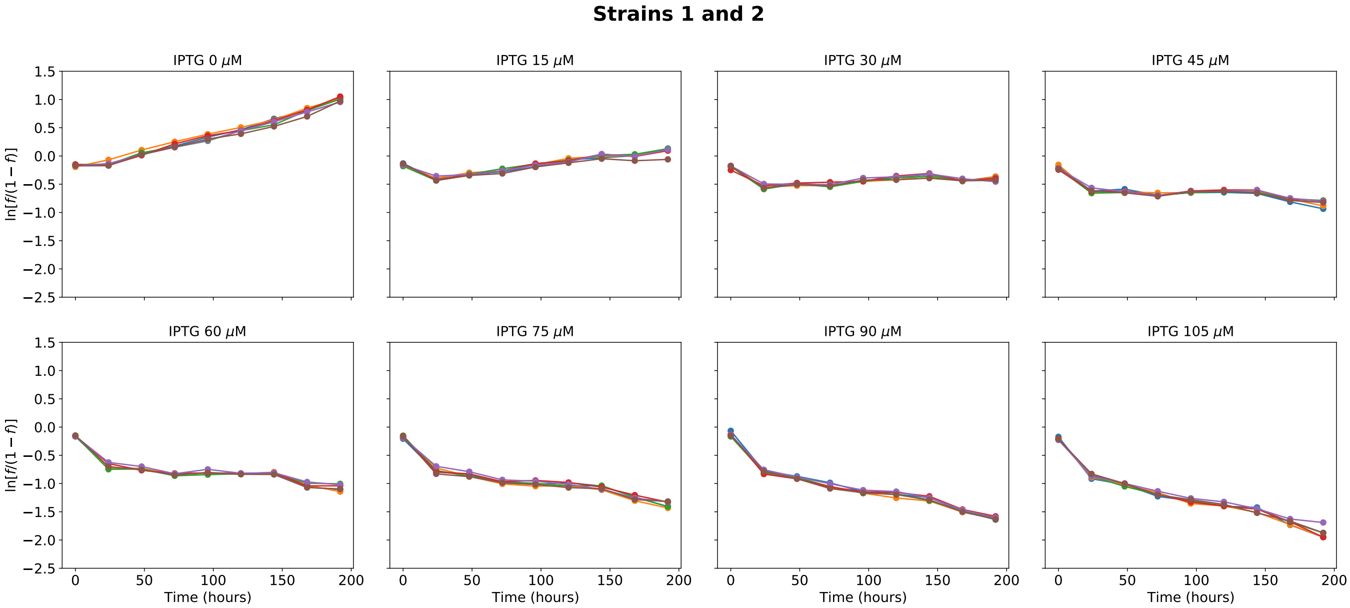


**FIG. S.2:** Time series of the values of $\ln f/(1-f)$ (with $f = m_1/(m_1+m_2)$ relative to the competition assay between strains 1 and 2. Each curve represent one of the six replicas of the treatment. The selective advantage $\mathcal{S}$ has been computed by fitting each of these curves with a linear function between day 1 and day 6 (i.e., between 24h and 144h).

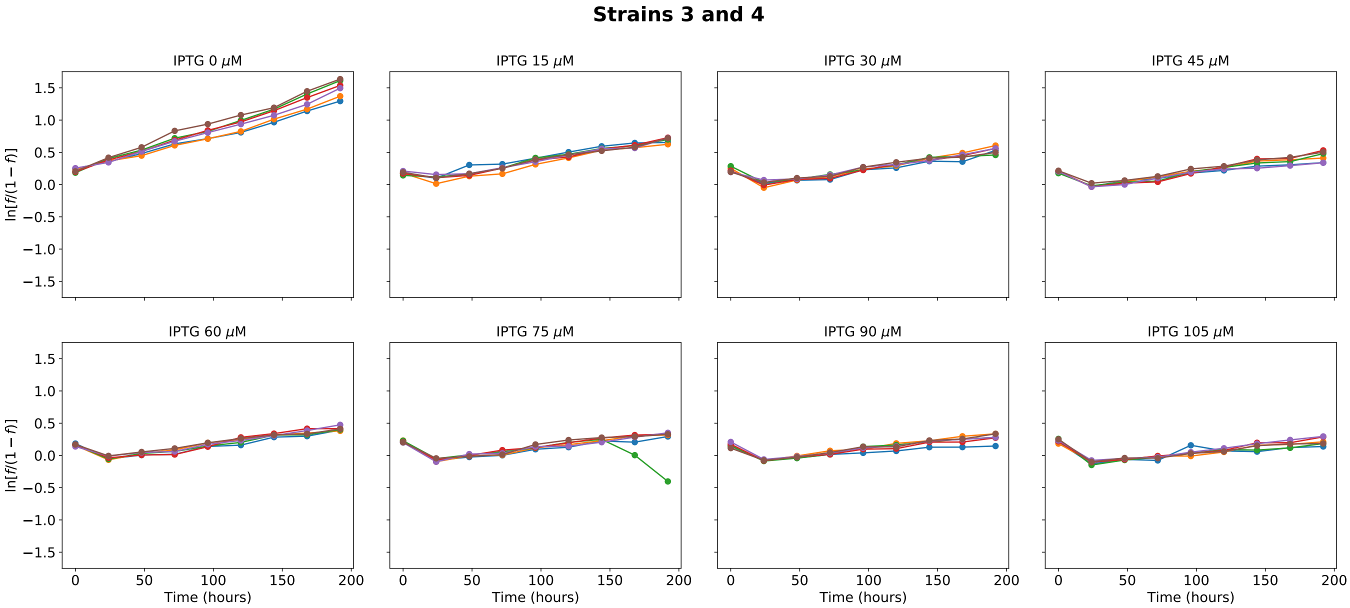


**FIG. S.3:** Time series of the values of $\ln f/(1-f)$ (with $f = m_3/(m_3+m_4)$ relative to the competition assay between strains 3 and 4. Each curve represent one of the six replicas of the treatment. The selective advantage $\mathcal{S}$ has been computed by fitting each of these curves with a linear function between day 1 and day 6 (i.e., between 24h and 144h).

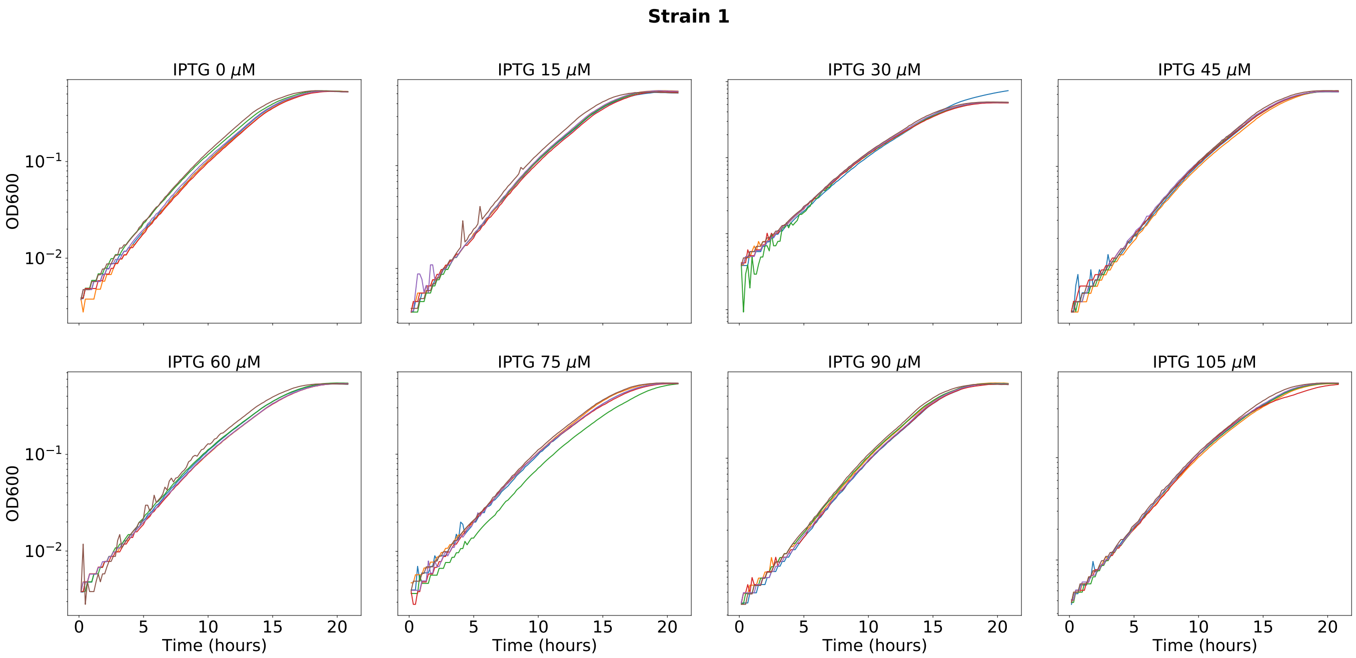


**FIG. S.4:** OD600 curves of strain 1 for different IPTG concentrations. These measures were obtained by preparing the strain with the same experimental protocol shown in the Methods section.

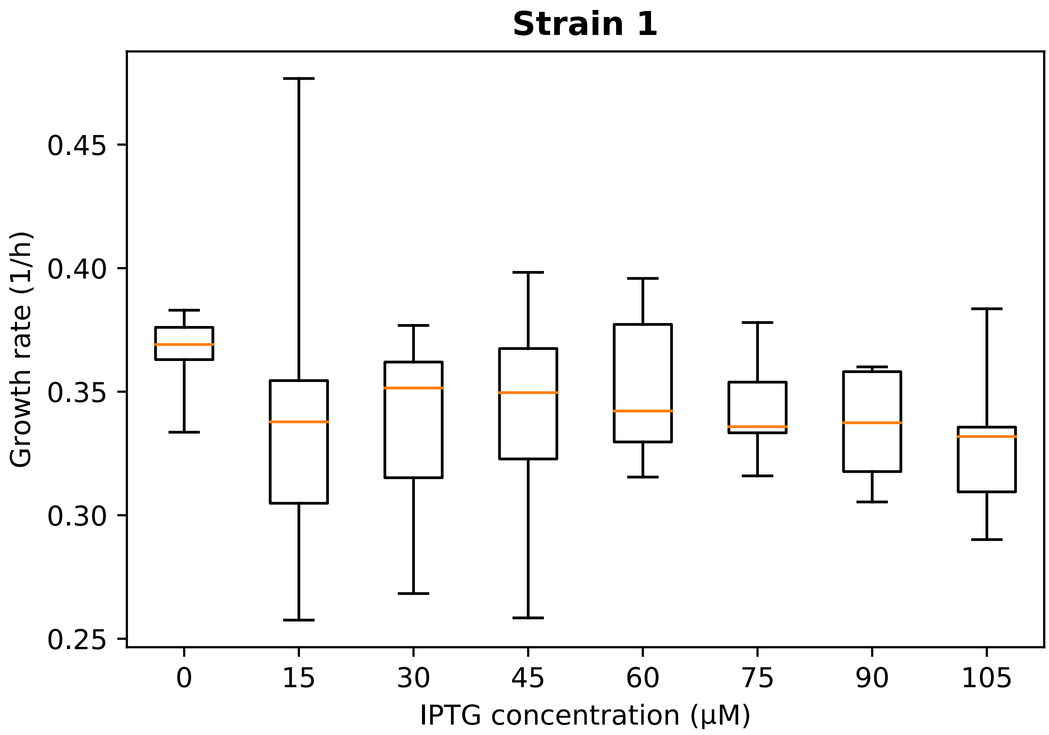


**FIG. S.5:** Boxplot of the growth rates of strain 1, obtained by fitting the curves in Figure S.4.

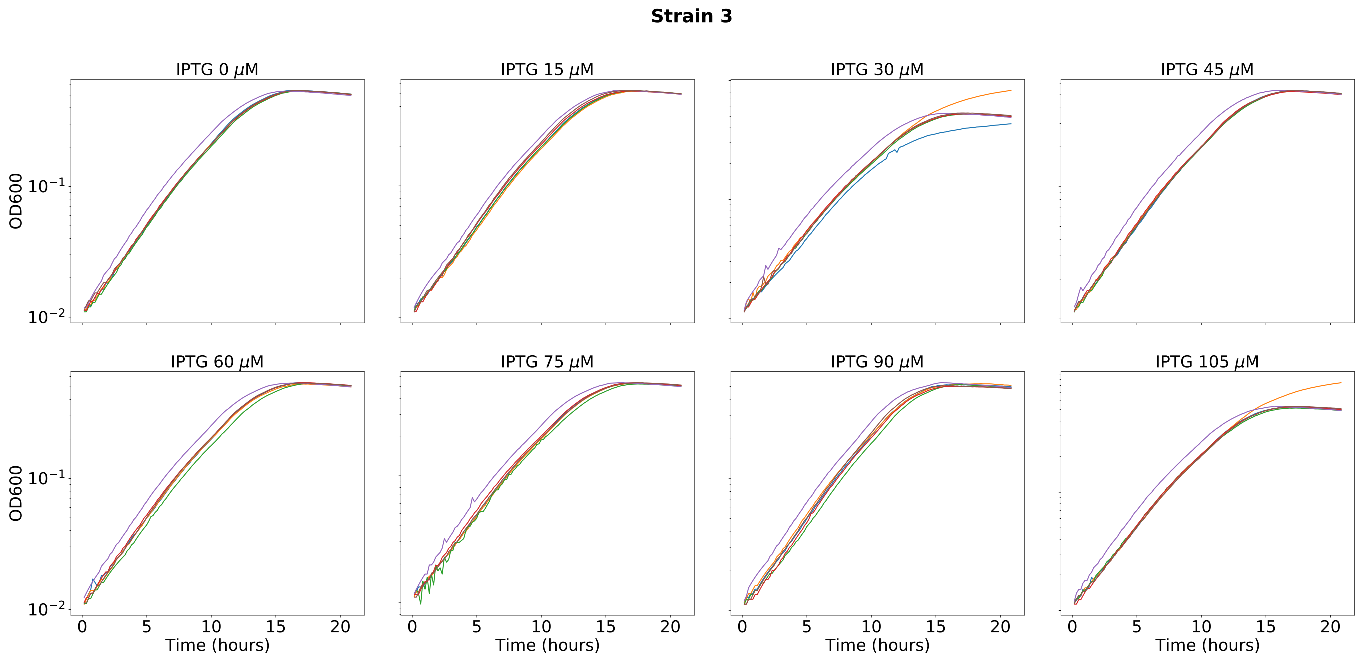


**FIG. S.6:** OD600 curves of strain 3 for different IPTG concentrations. These measures were obtained by preparing the strain with the same experimental protocol shown in the Methods section.

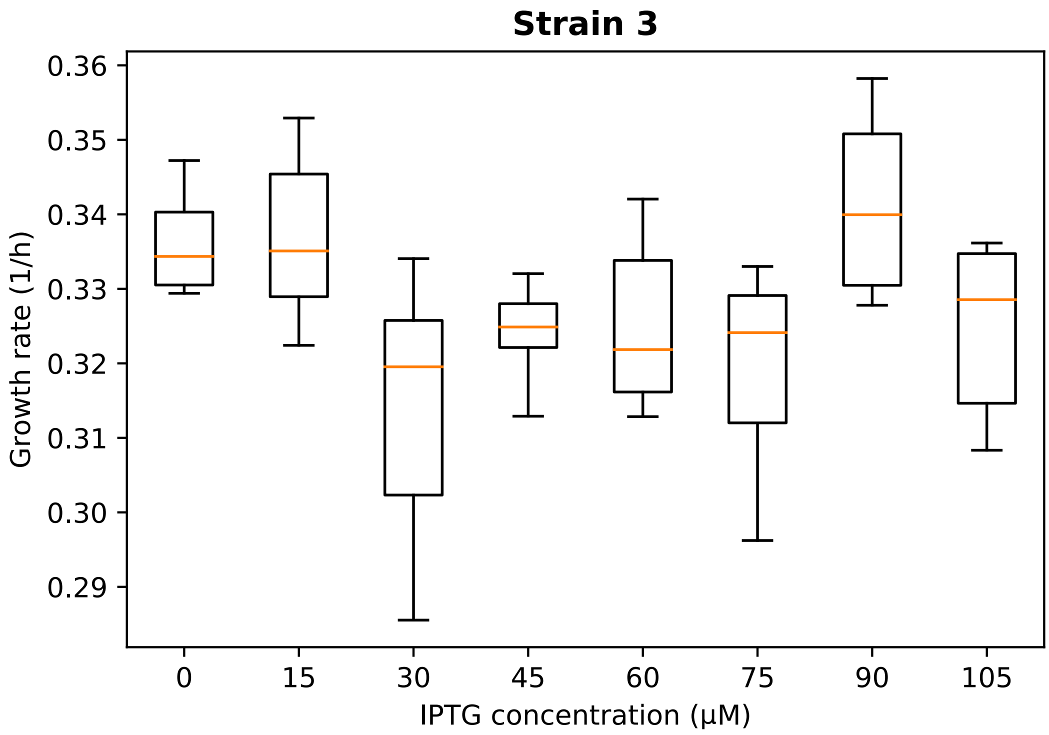


**FIG. S.7:** Boxplot of the growth rates of strain 3, obtained by fitting the curves in Figure S.6.

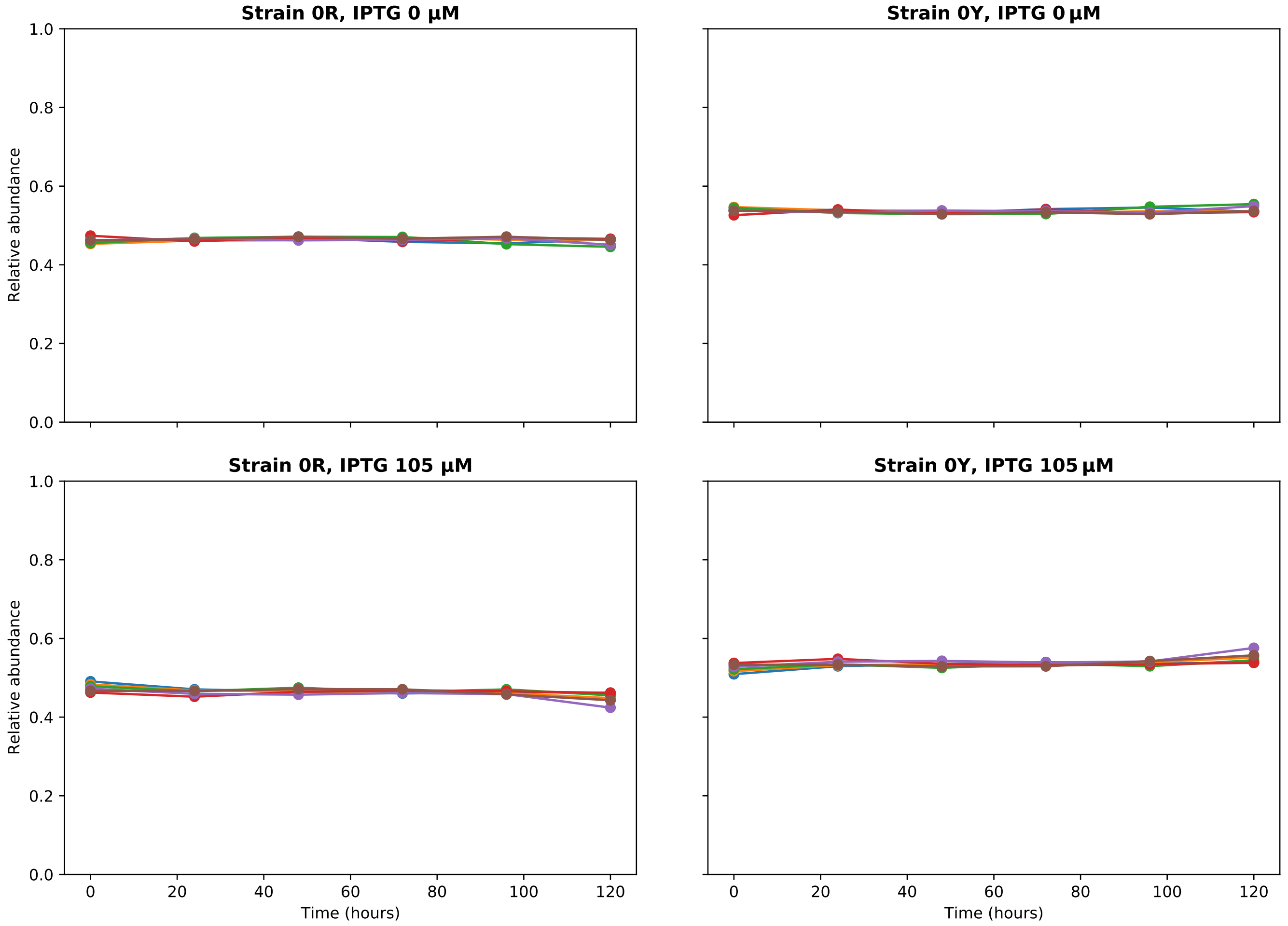


**FIG. S.8:** Competition assays between strains 0R and 0Y. These assays were done with the same experimental protocol shown in the Methods section (without adding ampicillin to the culture medium since both strains do not carry any plasmid and therefore are not ampicillin resistant).

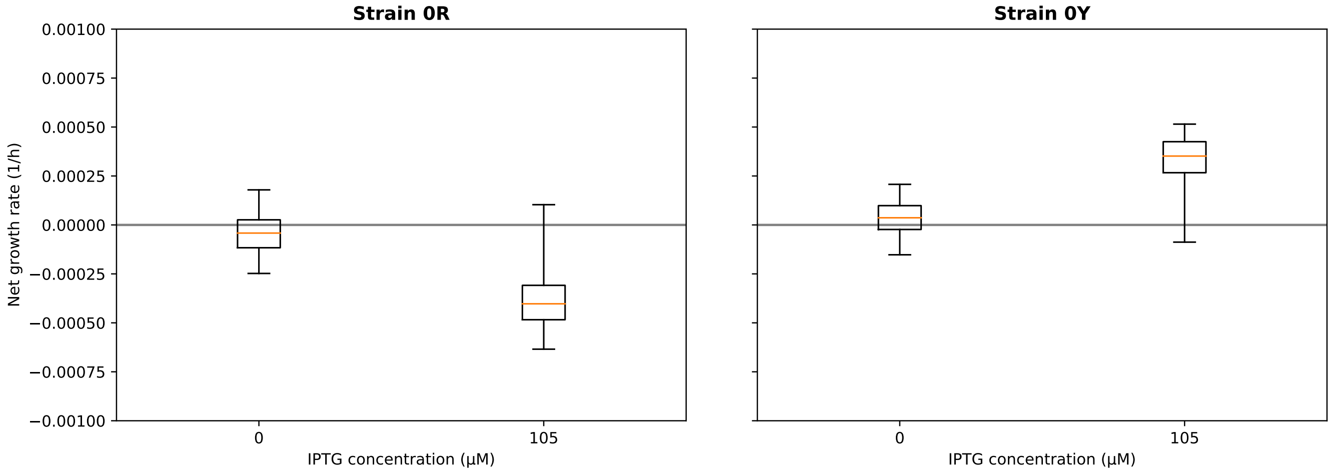


**FIG. S.9:** Boxplot of the growth rates of strains 0R and 0Y, computed by fitting the time series shown in Figure S.8.

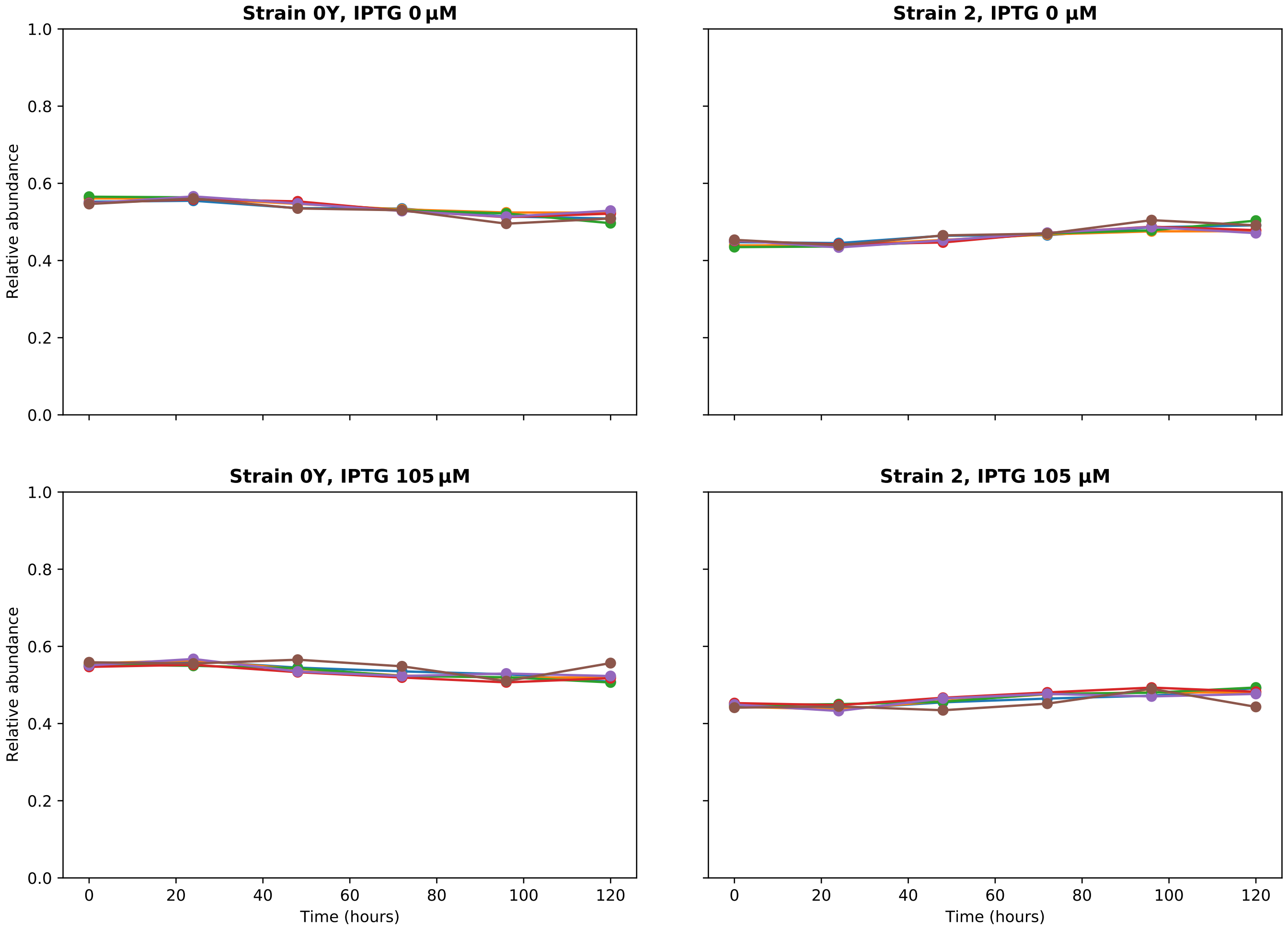


**FIG. S.10:** Competition assays between strains 0Y and 2. These assays were done with the same experimental protocol shown in the Methods section (without adding ampicillin to the culture medium since strain 0Y is not ampicillin resistant).

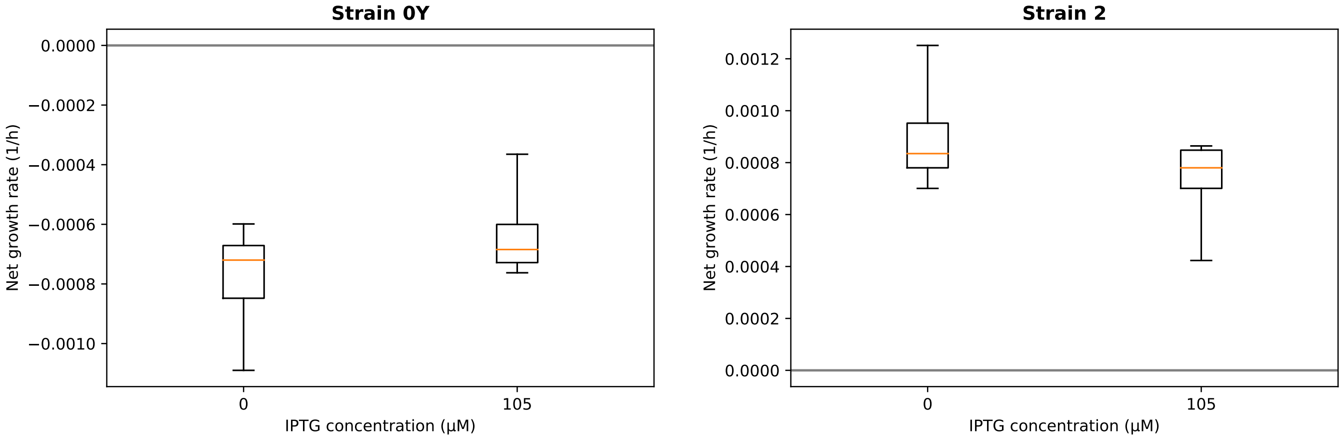


**FIG. S.11:** Boxplot of the growth rates of strains 0Y and 2, computed by fitting the time series shown in Figure S.10.

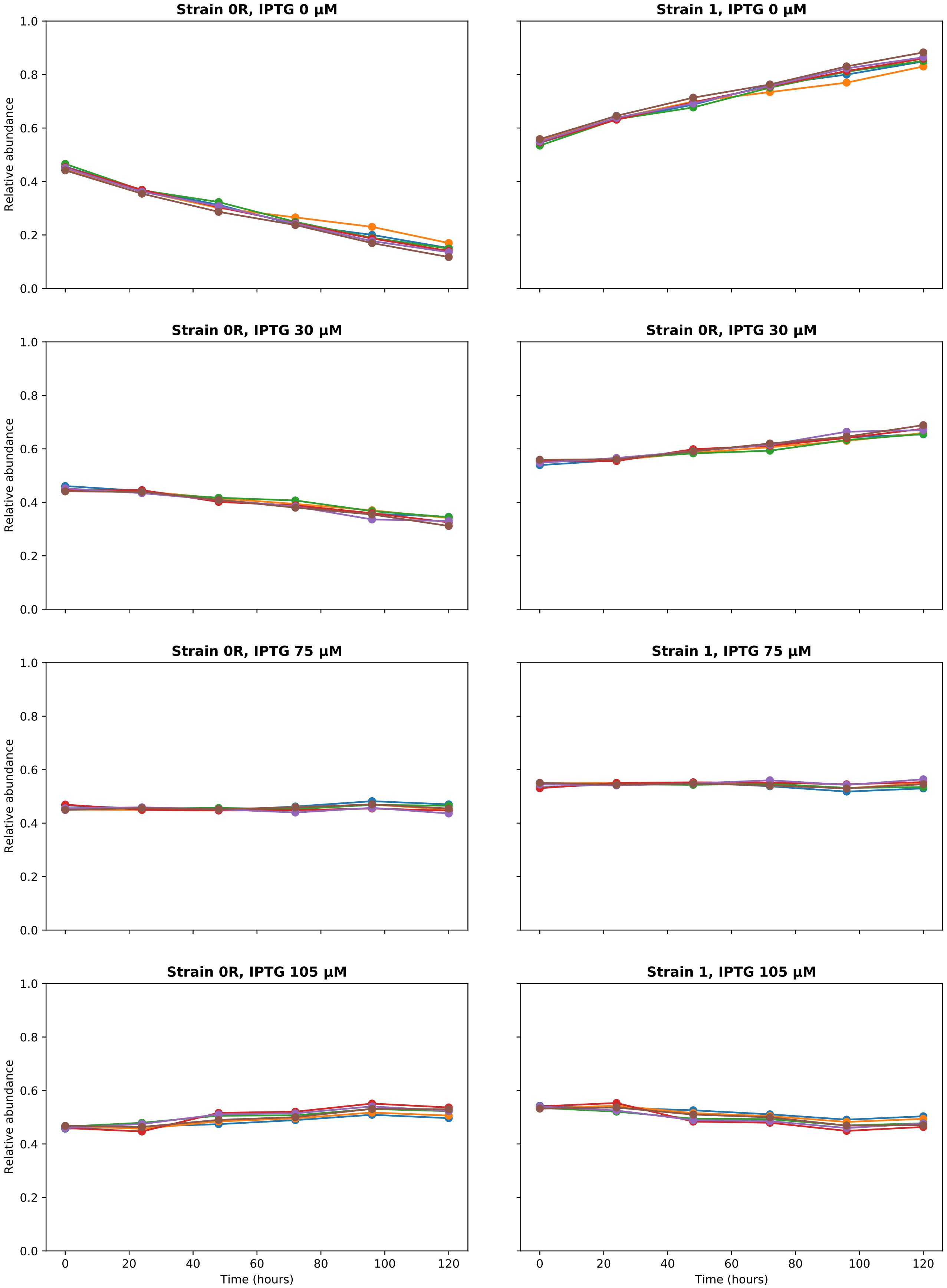


**FIG. S.12:** Competition assays between strains 0R and 1. These assays were done with the same experimental protocol shown in the Methods section (without adding ampicillin to the culture medium since strain 0R is not ampicillin resistant).

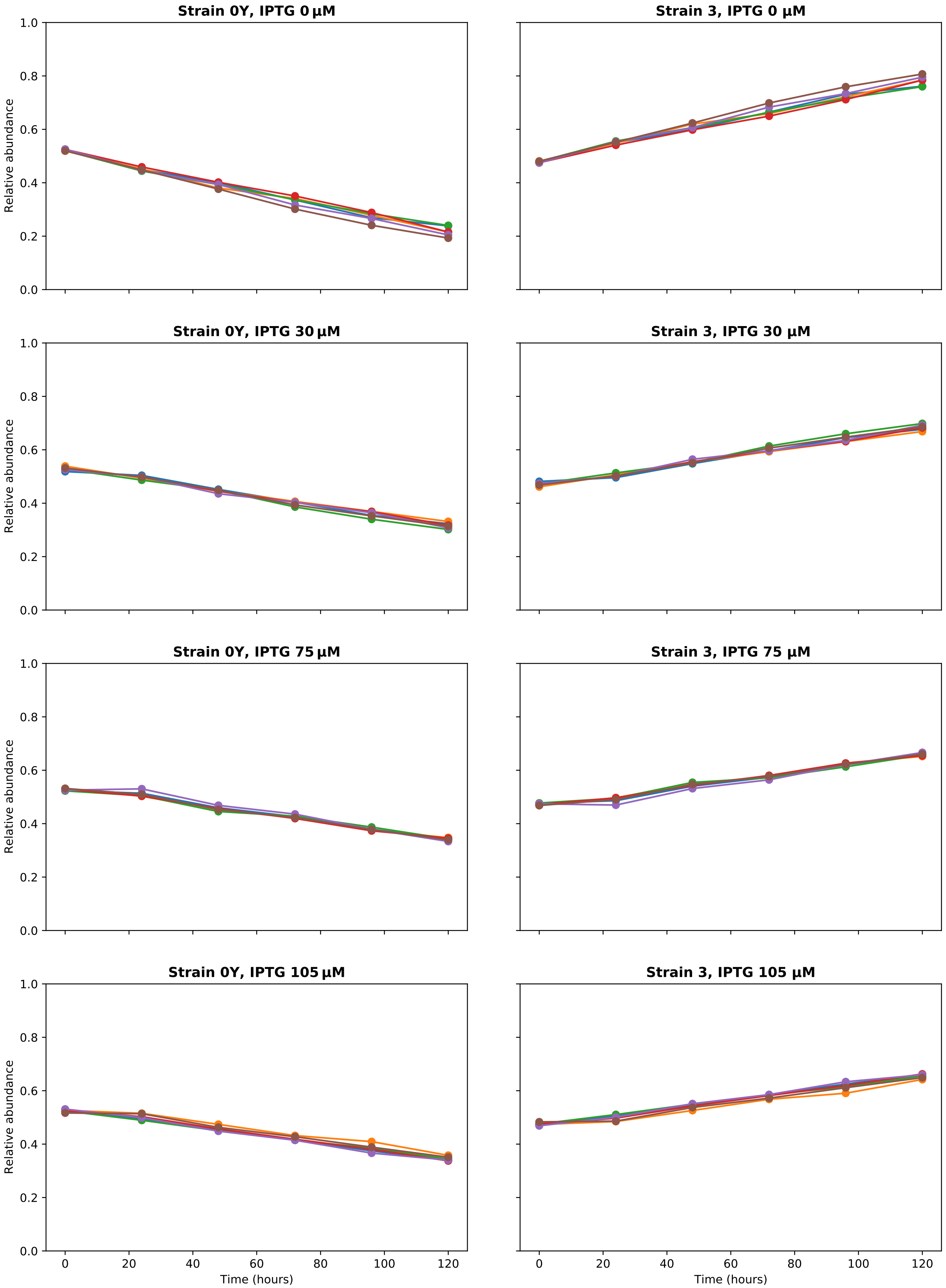


**FIG. S.13:** Competition assays between strains 0Y and 3. These assays were done with the same experimental protocol shown in the Methods section (without adding ampicillin to the culture medium since strain 0Y is not ampicillin resistant).

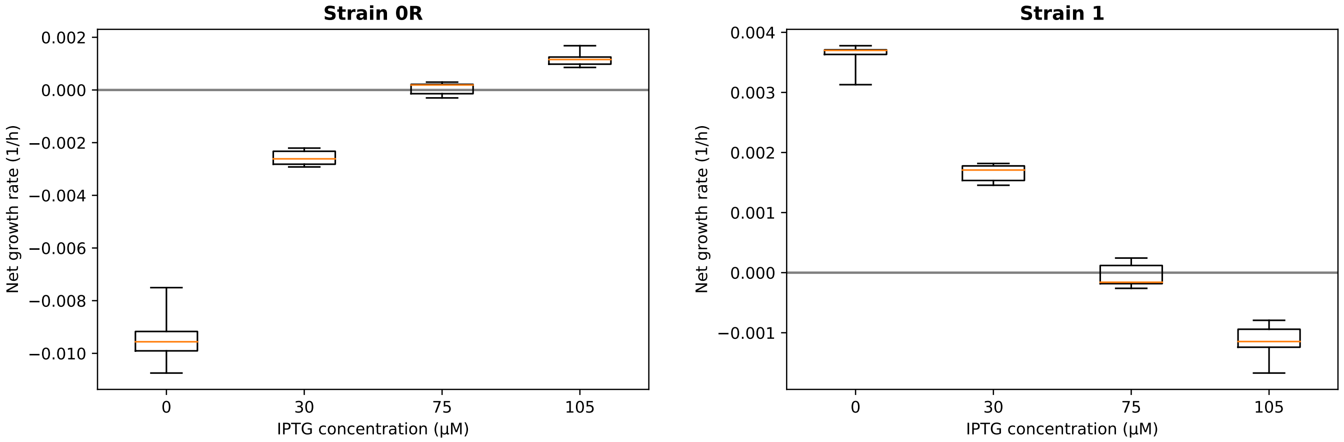


**FIG. S.14:** Boxplot of the growth rates of strains 0R and 1, computed by fitting the time series shown in Figure S.12. As we can see, in absence of IPTG strain 1 has a fitness advantage over 0R, which is reduced as the concentration of IPTG increases.

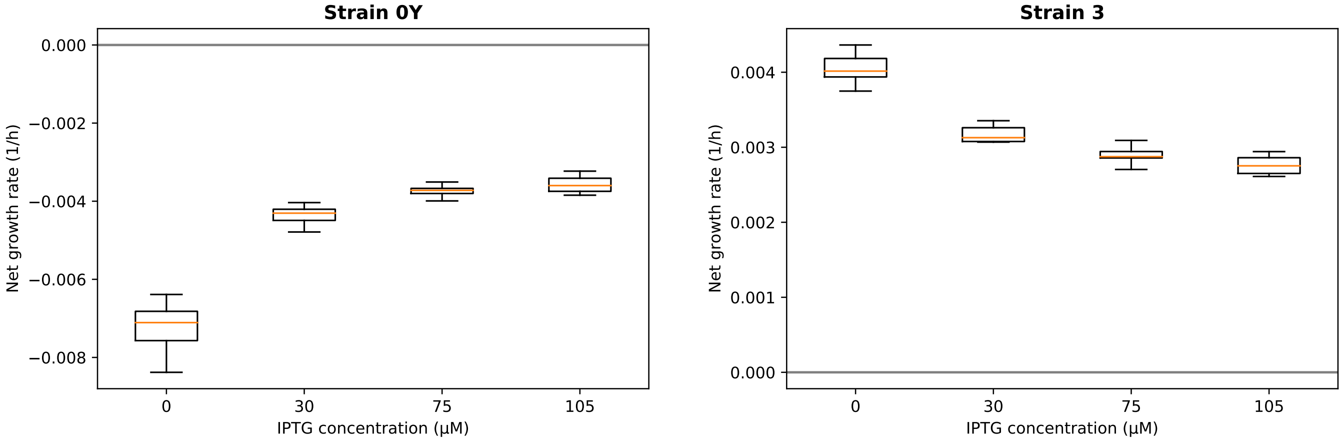


**FIG. S.15:** Boxplot of the growth rates of strains 0Y and 3, computed by fitting the time series shown in Figure S.13. As we can see, strain 3 always has a fitness advantage over 0Y.

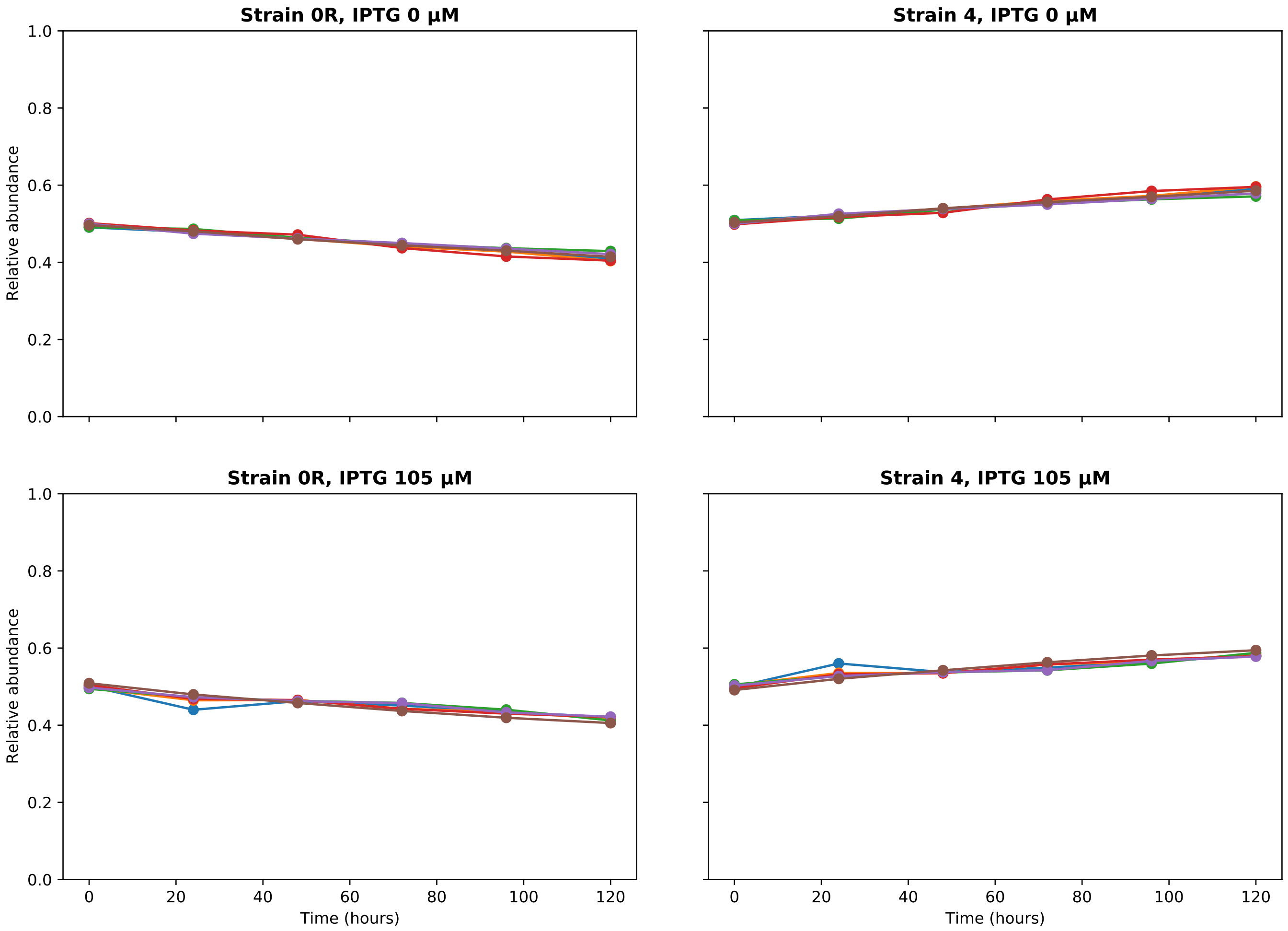


**FIG. S.16:** Competition assays between strains 0R and 4. These assays were done with the same experimental protocol shown in the Methods section (without adding ampicillin to the culture medium since strain 0R is not ampicillin resistant). As we can see, the plasmid containing ampicillin resistance carried by strain 4 confers a slight fitness advantage.

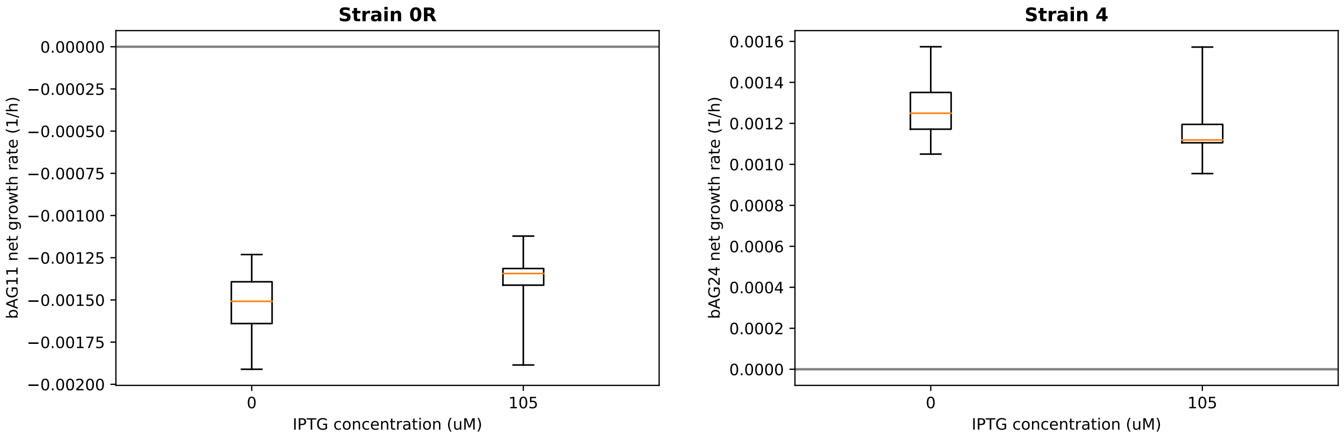


**FIG. S.17:** Boxplot of the growth rates of strains 0R and 4, computed by fitting the time series shown in Figure S.16

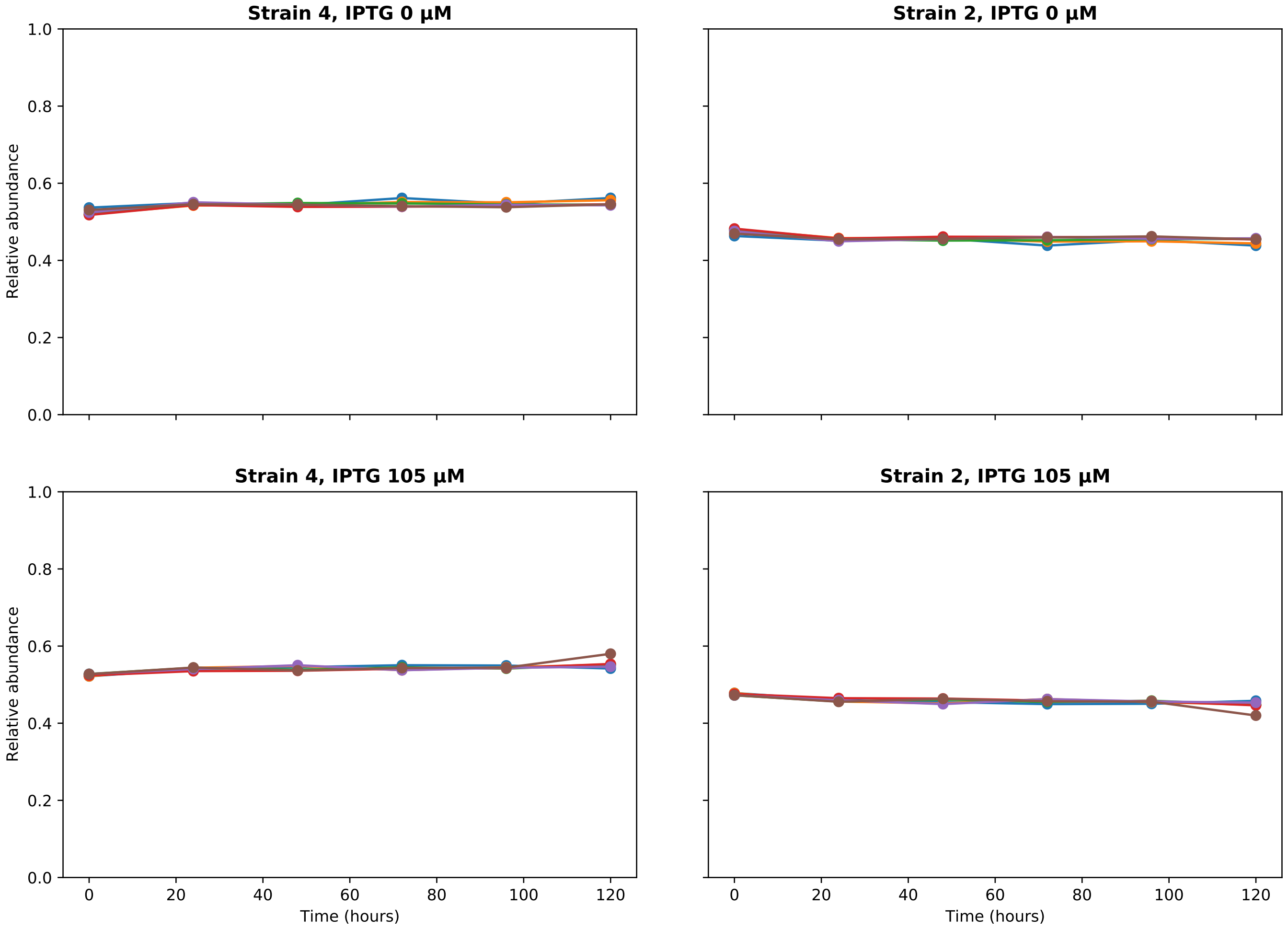


**FIG. S.18:** Competition assays between strains 4 and 2. These assays were done with the same experimental protocol shown in the Methods section.

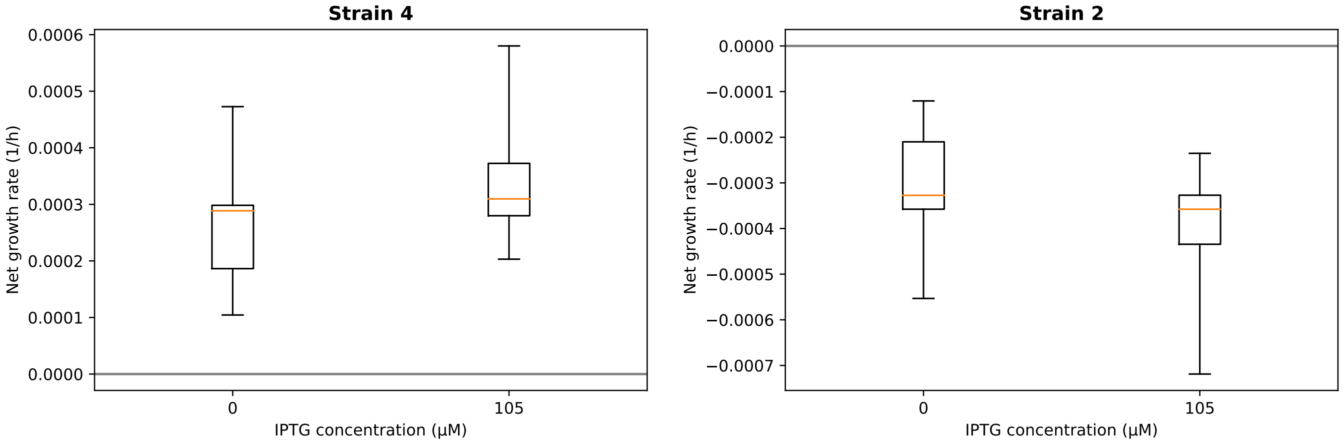


**FIG. S.19:** Boxplot of the growth rates of strains 4 and 2, computed by fitting the time series shown in Figure S.18

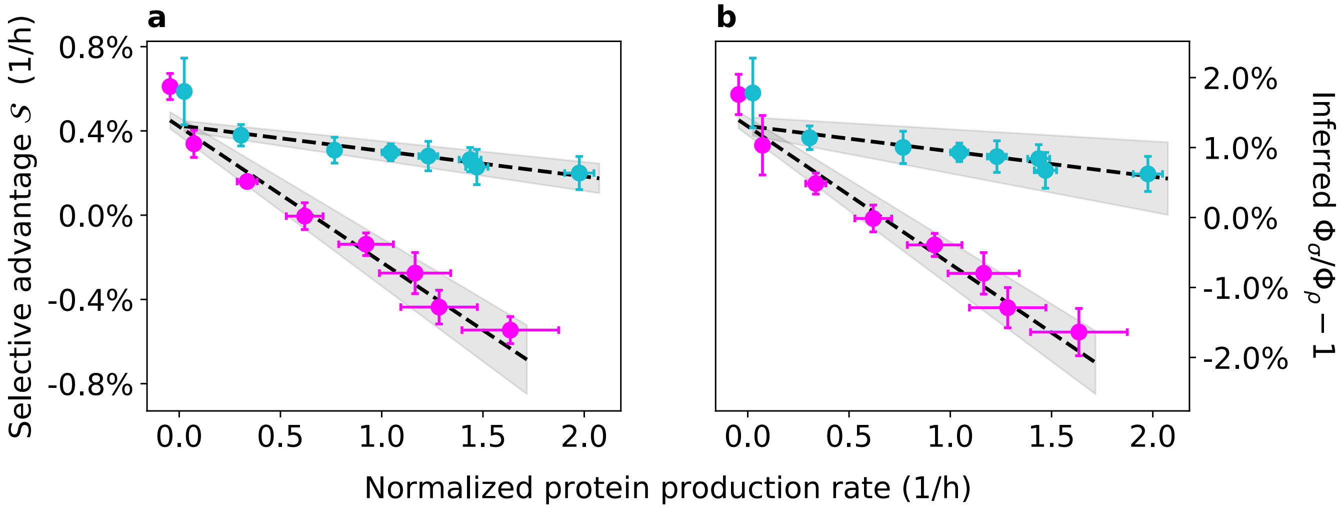


**FIG. S.20:** Results of the experiments (a) and inferred values of the ratios $\Phi_1/\Phi_2$ and $\Phi_3/\Phi_4$ (minus one), as shown in Figure 3 of the Main Text. In other words, magenta (cyan) points represent data of the experiment with strains 1 and 2 (3 and 4), error bars of the data points represent two standard deviations, and grey bands represent the 68% confidence interval of the linear fit). Differently from Figure 3, the linear fits shown here have been performed by including also the points at the lowest protein production rates (i.e., at 0 $\mu$M IPTG).
